# A nanobody-based strategy for rapid and scalable purification of native human protein complexes

**DOI:** 10.1101/2023.03.09.531980

**Authors:** Taylor Anthony Stevens, Giovani Pinton Tomaleri, Masami Hazu, Sophia Wei, Vy N. Nguyen, Charlene DeKalb, Rebecca M. Voorhees, Tino Pleiner

**Affiliations:** Division of Biology and Biological Engineering, California Institute of Technology, 1200 E. California Blvd., Pasadena, CA 91125, USA

## Abstract

Native isolation of proteins in high yield and purity is a major bottleneck for analysis of their three-dimensional structure, function, and interactome. Here, we present a streamlined workflow for the rapid production of proteins or protein complexes using lentiviral transduction of human suspension cells, combined with highly-specific nanobody-mediated purification and proteolytic elution. (1) First, generation of a plasmid coding for a protein of interest fused to an N- or C-terminal GFP or ALFA peptide tag is rapidly achieved using the lentiviral plasmid toolkit we have designed. (2) Human suspension cell lines stably expressing the tagged fusion protein can be generated in <5 days using lentiviral transduction. (3) Leveraging the picomolar affinity of the GFP and ALFA nanobodies for their respective tags, proteins expressed even at low levels can be specifically captured from the resulting cell lysate in a variety of conditions, including detergents and mild denaturants. (4) Finally, rapid and specific elution of tagged or untagged proteins under native conditions is achieved within minutes at 4°C using an engineered SUMO protease. We demonstrate the wide applicability of the method by purifying multiple challenging soluble and membrane protein complexes to high purity from human cells. Our strategy is also directly compatible with many widely used GFP expression plasmids, cell lines and transgenic model organisms; is faster than alternative approaches, requiring ∼8 days from cloning to purification; and results in substantially improved yields and purity.

## Introduction

A key prerequisite for many basic and pharmacological applications is the preparation of highly pure protein samples. Sample quality underlies the success of diverse experimental techniques including structural approaches (e.g. cryo-EM and X-ray crystallography), proteomics (e.g. interactome studies), and high-throughput studies of protein function (e.g. biophysical assays or *in vitro* drug screens).

Human proteins can be notoriously difficult to express and purify at the scale and purity needed for their structural and functional characterization. Their heterologous expression, e.g. in bacteria, frequently results in extensive degradation, insoluble aggregates or lack of essential post-translational modifications. These problems are exacerbated for multi-subunit protein complexes and membrane proteins, which may rely on mammalian specific factors for their biogenesis. Mammalian expression systems are thus in many cases the only alternative. However, culturing mammalian cells can be costly, making it essential to streamline both expression and protein purification workflows to maximize yield and purity, while saving valuable time and resources.

Here, we present a rapid and easily scalable workflow to purify soluble and membrane-spanning human proteins and protein complexes from human suspension cells for structural and functional analysis. Our workflow combines advances in both expression cell line generation^1^, as well as purification strategy^2^, and employs two highly versatile and complementary affinity tags – GFP and the small ALFA peptide tag^3^.

First, we describe how to rapidly generate stable polyclonal human suspension cell lines expressing a GFP- or ALFA-tagged protein of interest using lentiviral transduction. The use of lentivirus combines multiple advantages of both transient transfection and monoclonal cell line generation^1^. (1) Lentivirus is fast and cost-efficient, stable cell lines can be generated within <5 days using lentivirus produced from a single well of a 6-well plate. (2) It enables highly tunable expression, where transduction efficiencies can be titrated to either >80-90% to maximize yield, or to <30% to ensure a single copy of cDNA per cell. When optionally coupled with FACS, polyclonal cell lines can be generated that contain uniform expression across all cells. Finally (3), lentivirus ensures reproducible and scalable expression, because once generated, cell lines can be stored or expanded indefinitely. To streamline the cloning of suitable expression plasmids, we provide an extensive plasmid toolbox, an accompanying cloning guide, as well as general recommendations for GFP/ALFA tag placement.

Second, we present a nanobody-based strategy for the native isolation of human proteins that routinely results in higher yield and purity than canonical epitope tag-based approaches. This is achieved through the combination of high-affinity binding to anti-GFP or ALFA nanobodies with selective elution by an engineered SUMO protease (SENP^EuB^)^4^. SENP^EuB^ allows nearly 1,000-fold faster release of resin-bound proteins than TEV or 3C proteases, and elution can therefore be performed quickly under gentle conditions on ice. Rapid and selective elution of nanobody- captured proteins ensures high sample purity and preserves delicate protein complexes. We provide bacterial expression plasmids and protocols to inexpensively produce both nanobodies and protease in amounts sufficient for hundreds of purifications.

Through the combination of lentivirus-based cell line generation and nanobody-mediated affinity purification our protocol reduces the time and cost required to prepare high-quality protein samples. Together these strategies can therefore significantly accelerate the structural and functional analysis of otherwise difficult to express human protein complexes.

### Applications

One advantage of this protocol is that the nanobody-based purification strategy is highly flexible and modular. It can be used for purification of proteins or protein complexes under a variety of conditions and from lysates prepared from various eukaryotic or prokaryotic sources. In particular, utilizing the anti-GFP nanobody for protein capture can leverage the vast number of existing sources for GFP-tagged proteins (e.g. yeast, flies, worms and mice). Publicly available and commercial sources are enumerated in **Extended Data Figure 1**. Further, the nanobody•affinity tag pair can easily be exchanged to other natural, laboratory-evolved or computationally designed binding proteins, as well as protein-specific binders^5, 6^, making the system highly adaptable. A selection of potential options is listed in **Extended Data Figure 2**. We therefore envision that this strategy will be suitable for any structural or functional application requiring protein purification from a cell, extract, or tissue.

**Fig. 1.**
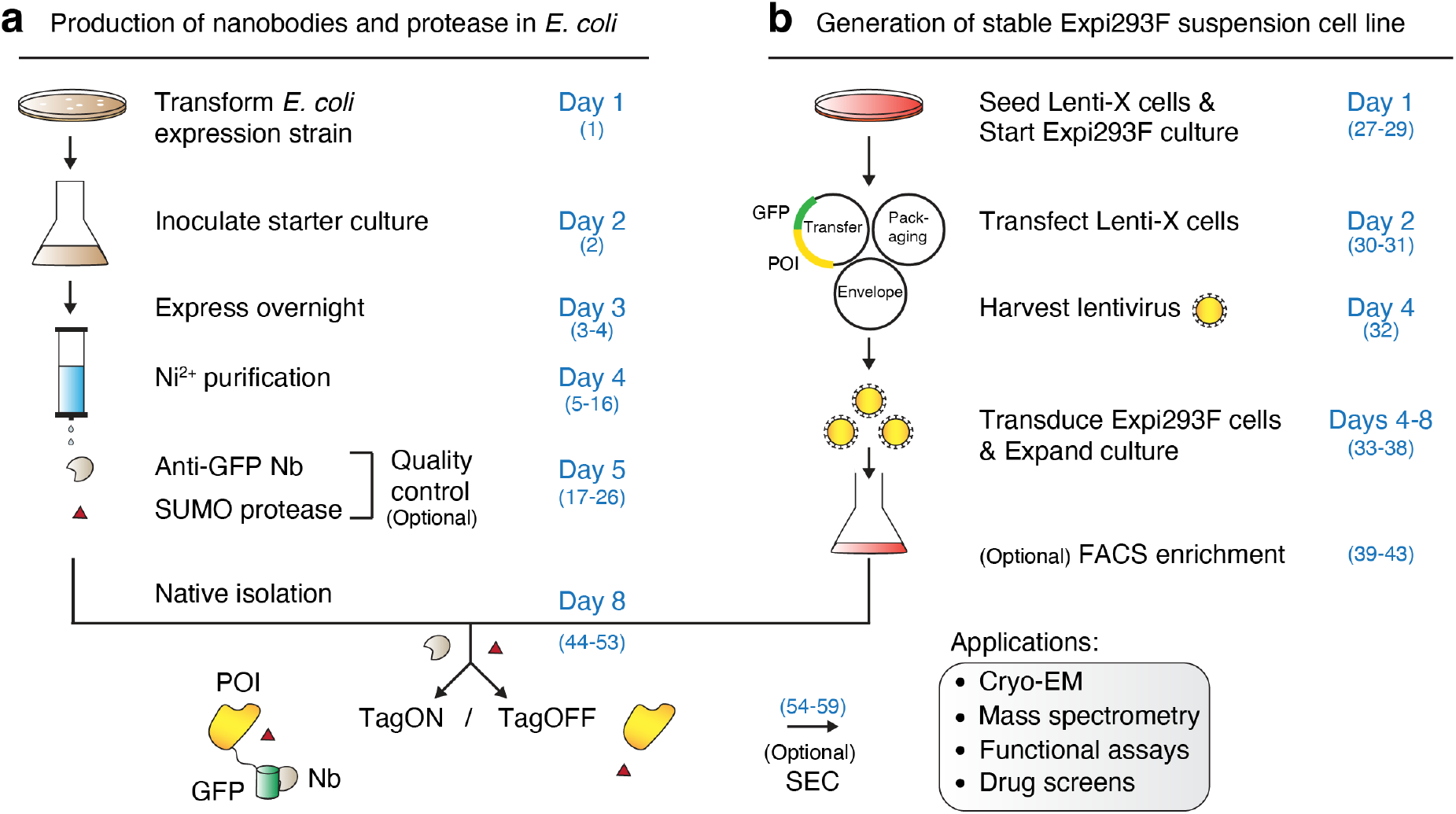
Schematic overview of the protocol. (**a**) Anti-GFP/ALFA nanobody (Nb)-based capture agents and SUMO protease are separately expressed in *E. coli* and purified via Ni^2+^-chelate affinity chromatography. (**b**) Lentivirus encoding a GFP/ALFA-tagged protein of interest (POI) is generated and used to transduce human Expi293F suspension cells. The resulting stable cell line is expanded and either used directly for first small-scale purification trials or optionally first sorted using fluorescence-activated cell sorting (FACS). In its fastest format our protocol can be completed in only 8 days, going from DNA prep to purified protein. Individual protocol step numbers are listed in parentheses. SEC = Size-exclusion chromatography.

**Fig. 2.**
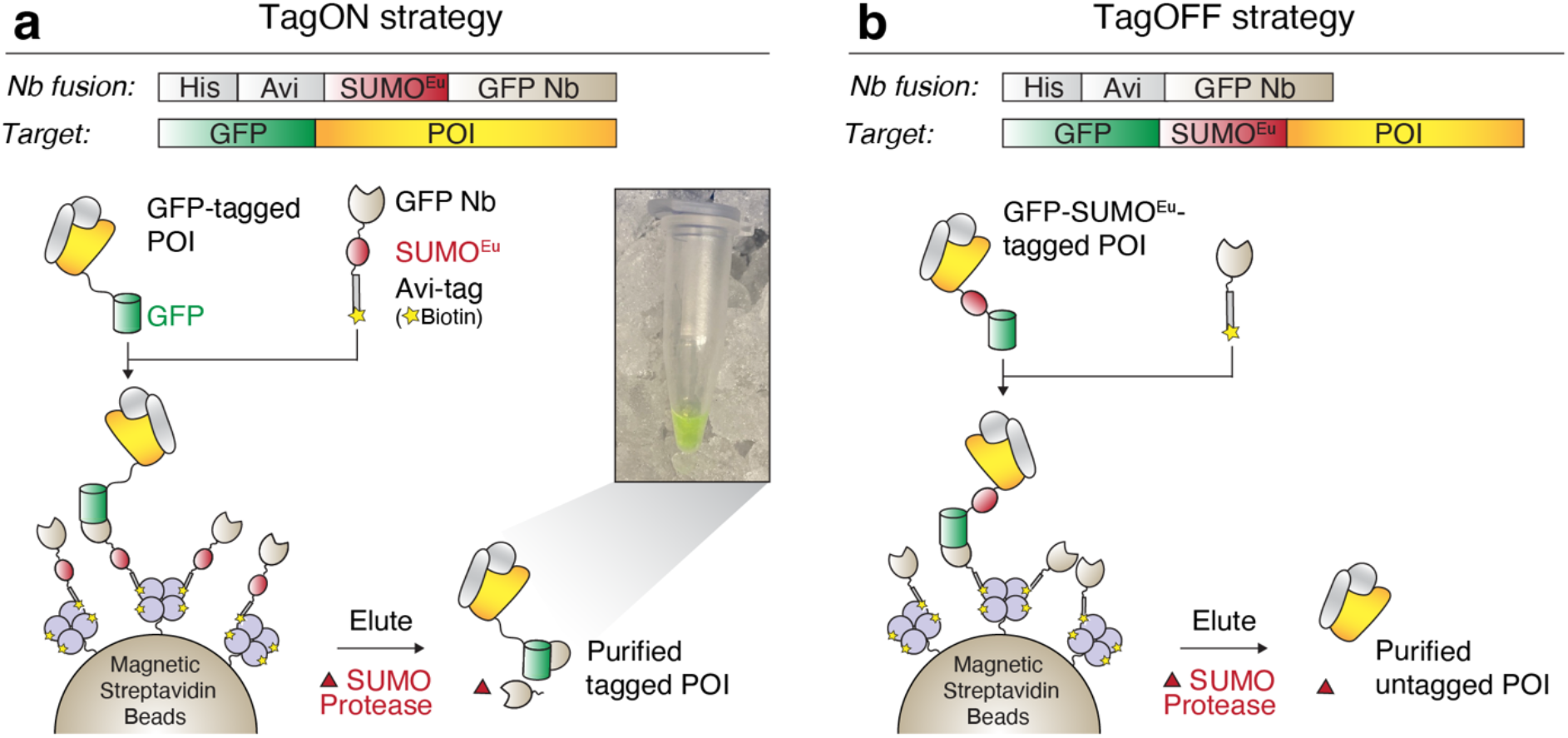
Native isolation of GFP/ALFA-fused proteins in tagged (TagON) or untagged form (TagOFF). (**a**) Schematic of the TagON strategy. A biotinylated SUMO^Eu^-fused anti-GFP nanobody (Nb) is immobilized onto magnetic streptavidin beads. Nb-decorated beads are then incubated with cell lysate containing an expressed GFP- tagged protein-of-interest (POI). Following wash steps, GFP-tagged POI and bound interaction partners (grey) are rapidly eluted by cleavage with the engineered SUMO protease SENP^EuB^. The inset depicts the result of a high-yield TagON purification (GFP-AE2, see Fig. 3c). (**b**) Schematic of the TagOFF strategy. SUMO^Eu^ is placed between GFP and POI and the POI is captured by a non-cleavable nanobody. This allows scarless tag-free elution of the POI by SENP^EuB^.

One particularly powerful application of this protocol is generating samples for structural analysis, which requires large scale production of highly pure samples^2, 7^. For example, we have used this strategy to purify the nine-subunit human ER membrane protein complex (EMC), for structure determination using single particle cryo-EM^2^. For the EMC, and many protein complexes, we have found that expression of a single tagged subunit results in its incorporation into the intact complex in place of the endogenous subunit. Any excess unassembled subunits are typically degraded by the ubiquitin-proteasome pathway, as has been previously described^8, 9^. Therefore, by exploiting endogenous protein quality control machinery in human cells, it is often possible to introduce a tag into a protein complex without expressing multiple subunits or first generating knock out cell lines.

Similarly, our protocol can be used to generate highly pure samples for functional assays (e.g. to characterize a protein’s enzymatic activity, stability or DNA/RNA/lipid-binding properties). We recently used it to isolate the 33 kDa mitochondrial outer membrane insertase MTCH2 to reconstitute its protein-insertion activity *in vitro*^10^.

Further, because of the high specificity, efficient capture, and rapid elution, this protocol is also well-suited for analysis of physical interaction partners by mass spectrometry.

### Overview of the procedure

Our protocol comprises the following steps that can be carried out in parallel. Additional details are outlined in the experimental design section (**Fig. 1**):

(i) Cloning of the protein of interest into a lentiviral transfer plasmid as a GFP- or ALFA tag fusion (for ease of use a plasmid toolbox is provided).
(ii) Generation of lentivirus and transduction of human suspension cells to stably integrate the desired open reading frame.
(iii) Recombinant expression of anti-GFP or ALFA nanobodies and protease in *E. coli* followed by Ni^2+^-chelate affinity purification.
(iv) Nanobody-mediated purification from the expanded suspension cell culture.

### Two different purification strategies – TagON and TagOFF

We present two different strategies to purify a GFP- or ALFA-tagged protein in either a tagged (TagON) or scarless untagged form (TagOFF) (**Fig. 2**). Both strategies employ distinct biotinylated anti-GFP or ALFA nanobody fusion proteins that are immobilized onto streptavidin beads to capture GFP- or ALFA-tagged proteins from cell lysate.

In the TagON strategy (**Fig. 2a**), a SUMO^Eu^ module is inserted between the biotin acceptor peptide (Avi) tag and nanobody, so that SENP^EuB^ cleavage releases the nanobody along with its bound protein. The eluted protein therefore retains the fluorescent GFP tag, which may be useful for detection during downstream applications such as Fluorescence detection size exclusion chromatography (FSEC)^11^. In the TagOFF strategy (**Fig. 2b**), a SUMO^Eu^ module is placed in between GFP/ALFA tag and the protein so that cleavage by SENP^EuB^ only releases the protein, while both nanobody and tag are retained on the resin. This strategy is useful if the presence of the tag interferes with downstream applications.

### Purification of soluble and membrane-bound multi-subunit protein complexes

To demonstrate the utility of both TagON and TagOFF, we used the nine-subunit EMC as a model substrate. GFP- or ALFA-tagged EMC2 or EMC5 are incorporated into the EMC in place of their respective endogenous subunits, allowing the intact complex to be isolated in high purity (**Fig. 3a- b**). Using the TagOFF strategy, capture of GFP-SUMO^Eu^-EMC2 with a non-cleavable anti-GFP nanobody allowed native isolation of completely untagged EMC following SENP^EuB^ cleavage, while the affinity tag and nanobody were retained on the beads.

**Fig. 3.**
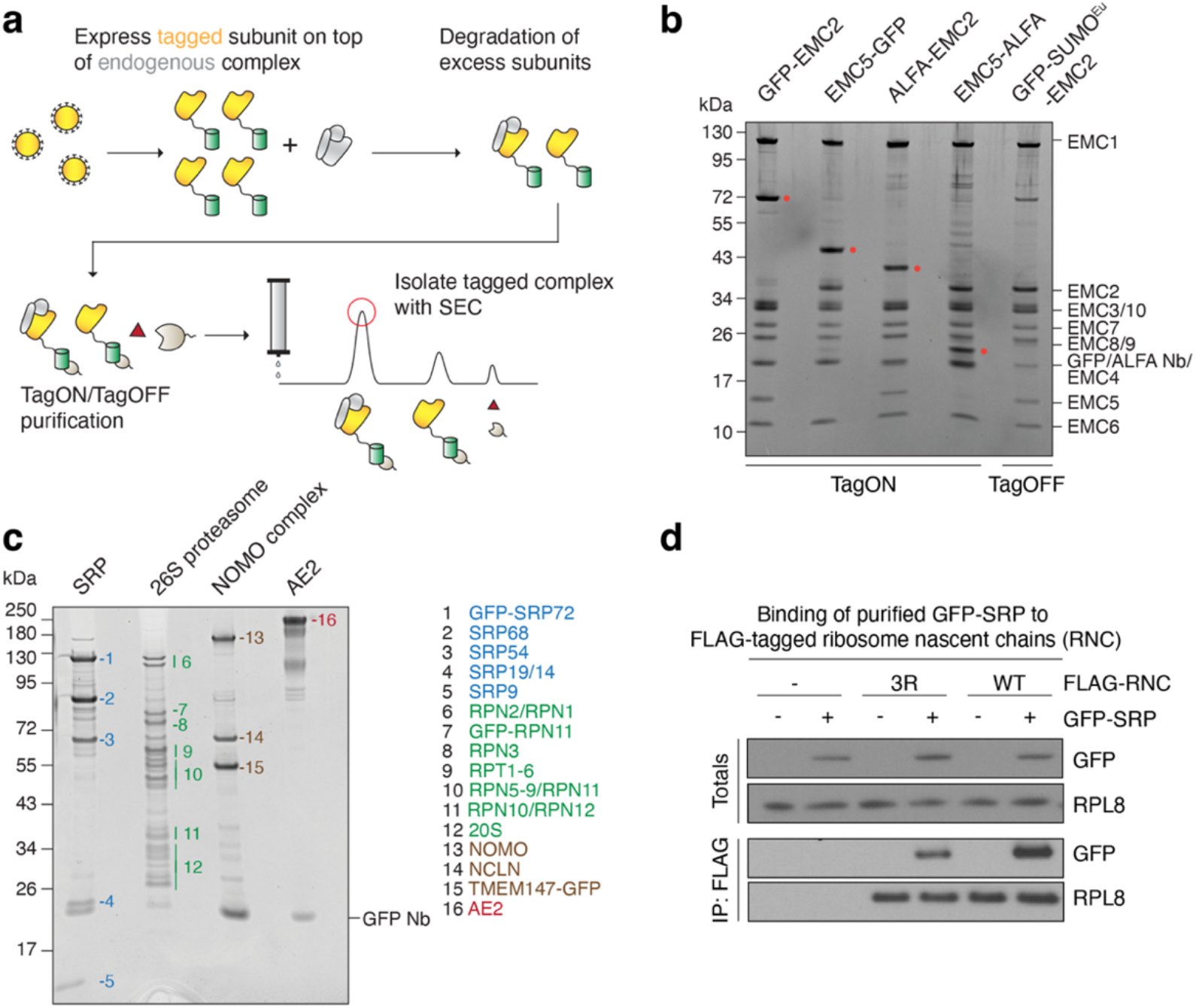
Purification of soluble and membrane protein complexes from human suspension cells. (**a)** Ectopically expressed tagged subunits of a protein complex replace their endogenous untagged counterparts (grey) through proteasomal degradation of excess subunits. After purification, any remaining excess subunit and nanobody can be removed via size-exclusion chromatography (SEC). (**b**) Peak fractions of SEC runs of the ER membrane protein complex (EMC) purified via GFP- or ALFA-tags fused to either EMC2 or EMC5 subunits using the TagON strategy. Following the TagOFF strategy, the GFP-SUMO^Eu^-EMC2 cell line allowed purification of completely untagged EMC. Tagged subunits are marked with a red dot. (**c**) SEC peaks of various samples purified via GFP-tags, including the signal recognition particle (SRP), 26S proteasome, NOMO-NCLN-TMEM147 complex, as well as SLC4A2/AE2. (**d**) Purified GFP-SRP is functional. Stalled ribosome nascent chain complexes (RNC) exposing either a wildtype (WT) or triple arginine mutant (3R) transferrin transmembrane domain (TMD) with 3x FLAG tag were produced by *in vitro* translation in rabbit reticulocyte extract supplemented with purified GFP-SRP complex where indicated. Total and FLAG-IP samples were analyzed by SDS-PAGE and Western blotting. GFP-SRP co-purified strongest with WT TMD RNCs as shown before for native SRP^12^.

As a proof-of-concept, we successfully isolated other challenging soluble and membrane protein complexes by tagging a single subunit (**Fig. 3c**). For example, GFP-SRP72 efficiently incorporated into the ribonucleoprotein signal recognition particle (SRP), which could be isolated in its native form. Purified SRP was fully functional and bound to stalled ribosome nascent chains exposing a transmembrane domain in a hydrophobicity sensitive manner as observed before for native SRP^12^ (**Fig. 3d**). Similarly, we used this strategy to isolate the entire 26S proteasome from cells expressing GFP-tagged RPN11, the ER-resident membrane protein complex NOMO-NCLN- TMEM147 via TMEM147-GFP, and the plasma membrane localized anion exchanger SLC4A2/AE2.

### Comparison with other methods

The advantages of this purification protocol over existing methods stem primarily from combining two innovations.

First, we achieve higher sample purity by using highly specific nanobodies and affinity beads. Our approach exploits the picomolar affinity between the nanobodies and their epitope tags to selectively capture tagged proteins from cell lysate. Additionally, both the nanobody fusion proteins and affinity resin are fully orthogonal to eukaryotic and prokaryotic host proteins. For example, the nanobody-decorated streptavidin beads are passivated by blocking any excess binding sites with free biotin or charged PEGylated biotin derivatives. Therefore, in contrast to the TwinStrep:Streptactin system, our strategy does not capture endogenous biotinylated proteins.

Second, rapid and selective protease elution of nanobody-captured proteins additionally ensures high sample purity and quality. In contrast, commonly used epitope tag-binding monoclonal antibodies (e.g. FLAG, HA, Myc, or V5), rely on native elution using excess epitope peptide. This is often highly inefficient and requires prolonged incubation at room temperature, which releases background binders and can cause aggregation of fragile proteins or dissociation of weaker binding partners. A side-by-side comparison shows that EMC isolated from an EMC5-GFP cell line using SENP^EuB^ cleavage is substantially purer than EMC isolated from an EMC5-3xFLAG cell line using traditional anti-FLAG purification (**Fig. 4a**). The combination of specific nanobodies, passivated affinity resin and protease elution therefore yields high final sample purity.

**Fig. 4.**
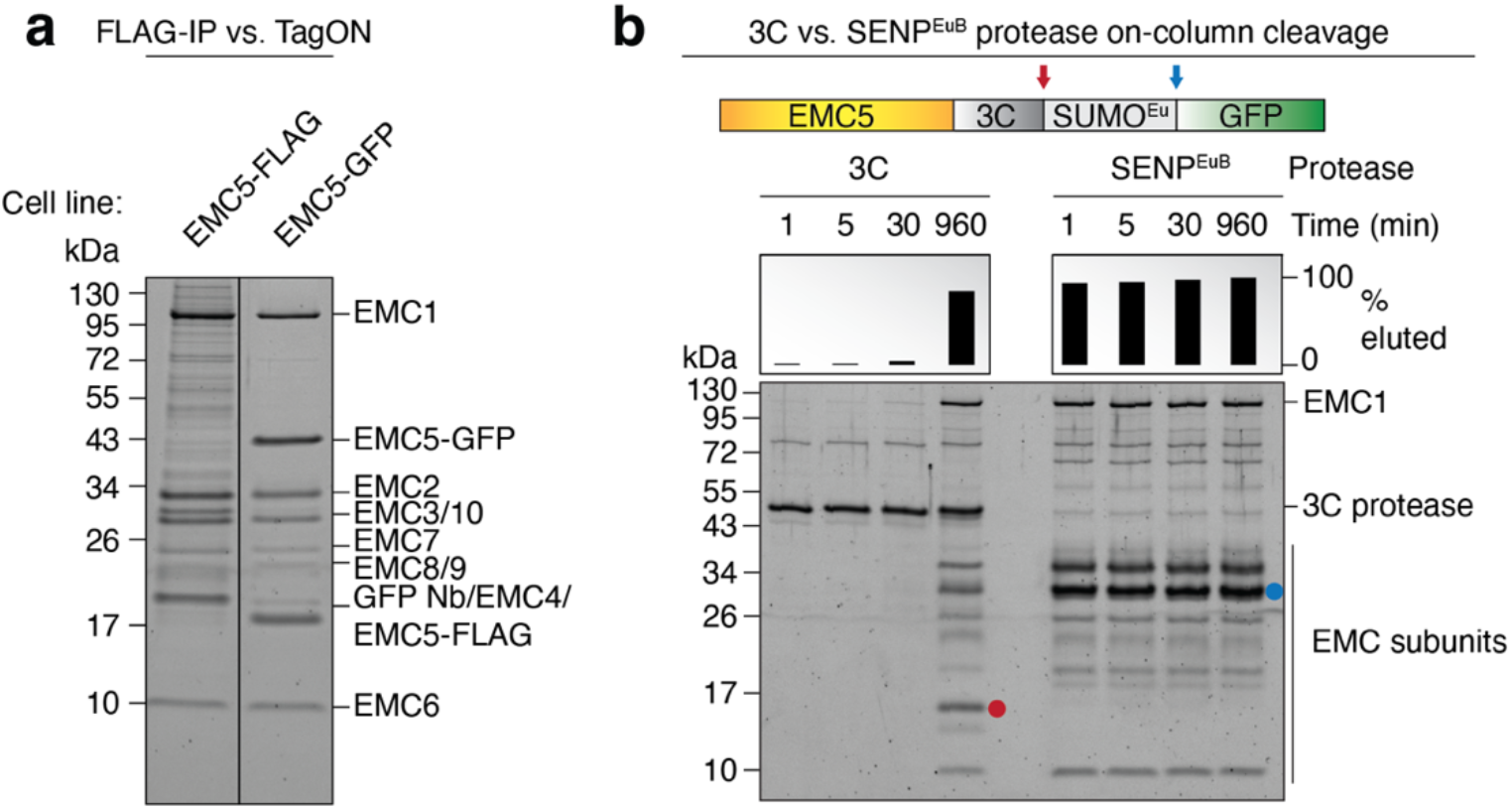
Comparison to other methods. (**a**) Comparison of FLAG and TagON purification of the ER membrane protein complex (EMC). Elutions from either EMC5-FLAG (using M2 FLAG affinity resin [Millipore-Sigma, USA]) or EMC5-GFP (using TagON) purifications were analyzed by SDS-PAGE and Sypro Ruby staining. (**b**) Comparison of 3C and SENP^EuB^ protease on-column cleavage efficiency. Purification of EMC via EMC5-3C-SUMO-GFP (using the TagOFF strategy). Cell lysate was incubated with magnetic Streptavidin beads, containing immobilized non-cleavable biotinylated anti-GFP nanobody (expressed from pTS117). After washing, beads were split in half and incubated either with 2 µM 3C or 250 nM SENP^EuB^ protease at 4°C for the indicated time frames. Under these conditions, SENP^EuB^ cleaves nearly 1,000-fold faster, allowing for rapid elution in just 1 min as opposed to lengthy overnight incubations typically required for elution with 3C or TEV protease. The products of 3C and SENP^EuB^ cleavage are marked with red and blue dots, respectively.

Protease cleavage is thus a powerful alternative over inefficient competitive peptide elution. However, common proteases like human rhinovirus (HRV) 3C protease or Tobacco etch virus (TEV) protease inefficiently cleave resin-bound proteins, especially on ice. Often large amounts of contaminating protease and lengthy overnight incubations (>13h) are needed to circumvent these issues. In contrast, low nanomolar concentrations of SENP^EuB^ are sufficient to gently release resin-bound proteins on ice within a few minutes (**Fig. 4b**).

Through both innovations, our protocol thus improves the signal-to-noise ratio of affinity purifications and better preserves sensitive protein complexes and their transient interaction partners. This yields less false interactors for affinity purification mass spectrometry (AP-MS) experiments and should also reduce contaminants that may interfere with functional experiments.

## Experimental design

### Optimal introduction of a tag on a protein subunit or complex

The first critical consideration for the purification of a multi-subunit protein complex is to choose a subunit to tag. We frequently found that ectopic expression of a single tagged subunit of a protein complex in human cells can result in the replacement of its endogenous counterpart. This strategy yields stoichiometric complexes in human cells when efficient cellular quality control mechanisms exist that degrade excess tagged and endogenous subunits of the chosen complex. Existing functional and structural data can often be used to inform subunit choice. For example, scaffold subunits of protein complexes typically contain multiple hydrophobic interfaces and are especially short-lived in the unassembled state, ensuring rapid elimination of excess subunits. Such subunits might thus represent prime candidates for first tagging trials^9^ and can be identified from mass spectrometry studies that analyzed proteome-wide degradation kinetics^13^. More stable subunits can accumulate after overexpression and may need to be removed by additional purification steps.

The next important consideration is to identify the optimal location of the affinity tag to minimize its interference with protein function. For protein complexes, the tag must be compatible with complex assembly and integrity. Compatibility can often be inferred from prior published studies or large-scale GFP-tagging efforts (**Extended Data Figure 1**), provided a careful analysis of localization and function was performed. Alternatively, existing structures or structural models of the proteins can be analyzed. A flexible unstructured terminus that is not part of a folded domain can usually tolerate a protein tag. Conversely, regions with high sequence conservation might indicate a functional requirement or binding site. In general, smaller peptide tags (like the ALFA tag) tend to be less disruptive than larger globular tags. The ALFA tag can also be placed internally into exposed loops and thus provides an alternative for proteins that cannot be tagged at either terminus.

If the presence of a tag will interfere with downstream applications, the TagOFF strategy should be used to completely remove the tag during purification. For this, the appropriate non-cleavable nanobody fusion protein needs to be generated and combined with a protein expression vector that contains a protease cleavage site. As described below, N-terminal tags can be removed completely (scarless), yet C-terminal tag removal will always generate a cleavage scar.

### Cloning of a lentiviral transfer plasmid encoding the tagged protein of interest

Once a tagging and purification strategy is identified, the coding sequence of the protein of interest needs to be cloned into a lentiviral transfer plasmid. We provide an extensive toolbox of 2^nd^ generation lentiviral transfer plasmids to facilitate fusion of a protein of interest to a GFP or ALFA tag. All plasmids are available via Addgene (see **Table 1** for IDs). Each tag is either N-terminal (pTS93-pTS102) or C-terminal (pTS103-pTS116) and either non-cleavable for TagON or cleavable for TagOFF purification. A detailed cloning guide is provided in the supplements (**Supplementary Data 1**).

**Table 1.**
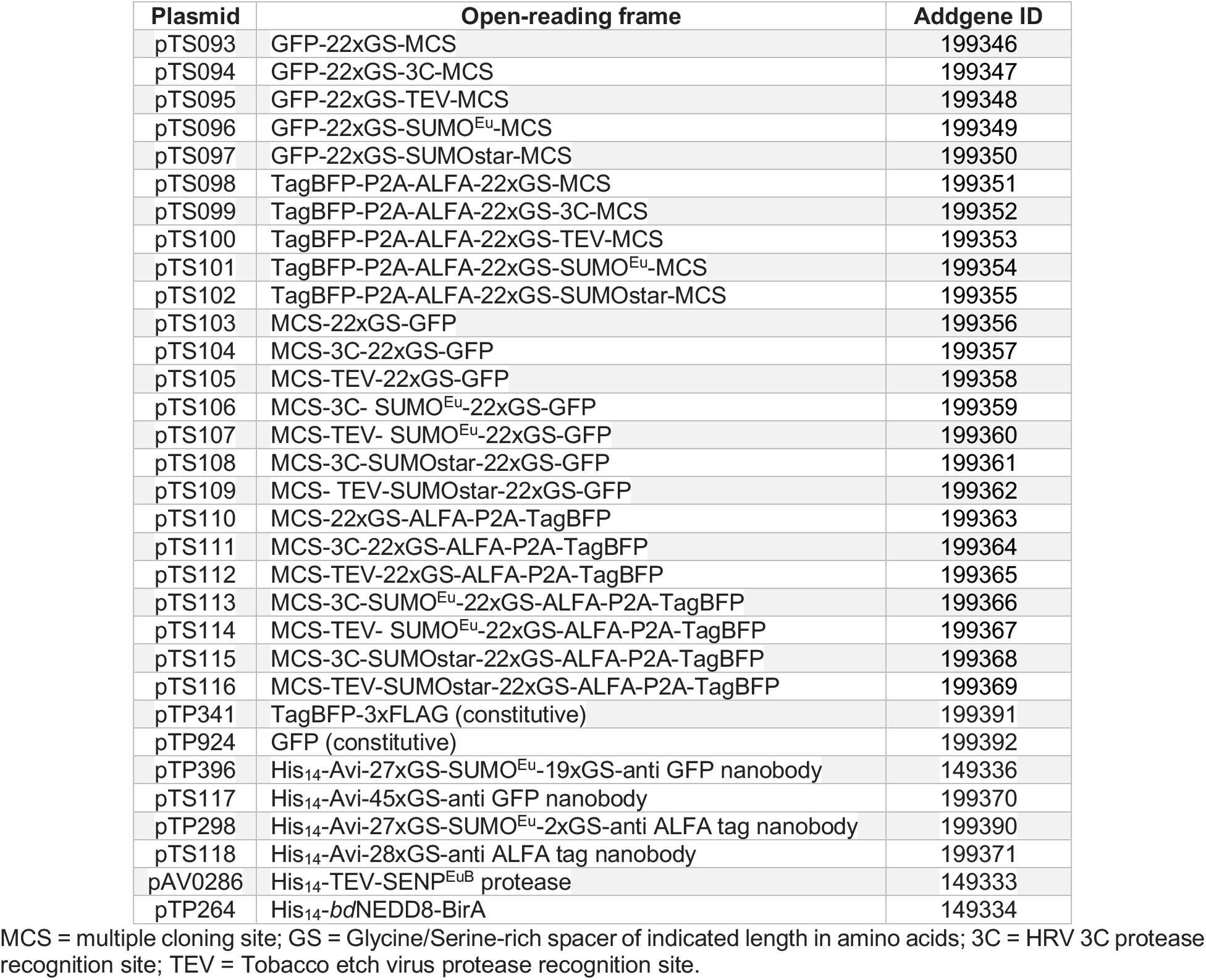
TagON and TagOFF-compatible lentiviral transfer plasmid toolbox and E. coli expression plasmids.

All constructs contain a CMV promoter fused to two Tet operator sequences (CMV-TetO2), which are Tet repressor (TetR) binding sites. The CMV-TetO2 promoter is thus doxycycline-inducible only in a TetR+ cell line, but will otherwise express constitutively.

We strongly recommend homemade SENP^EuB^ protease for tag removal, but we also provide plasmids encoding the cleavage sites of the commercially available 3C, TEV and SUMOstar proteases. Because these proteases all cleave at the C-terminus of their recognition site, N-terminal tags can thus be nearly completely removed. For C-terminal tags, however, this results in a cleavage scar. The scars of 3C and TEV protease (both 6 amino acids [aa]) are smaller than those of the SENP^EuB^ (96 aa) and SUMOstar proteases (99 aa). If desired these larger C-terminal scars can be removed after elution by an additional 3C or TEV protease cleavage step in solution. Our SUMO protease-cleavable C-terminal GFP or ALFA-tag encoding plasmids thus contain additional 3C or TEV cleavage sites between multiple cloning site and SUMO module.

One consideration in construct selection is that to maximize protein yield it can be beneficial to isolate successfully transduced cells via fluorescence-activated cell sorting (FACS) to obtain a homogenous population. While cells expressing GFP-tagged proteins can easily be sorted using GFP fluorescence, we additionally equipped expression vectors encoding ALFA-tagged proteins with a TagBFP expression cassette. TagBFP is separated from the ALFA-tagged protein using a porcine teschovirus P2A sequence, which mediates peptide bond skipping by the ribosome^14^ and thus results in the efficient synthesis of two separate proteins from a single mRNA – TagBFP and the ALFA-tagged protein.

### Generation of stable human suspension cell lines using lentiviral transduction

Lentiviral particles can be generated by co-transfecting adherent Lenti-X 293T cells with a 2^nd^ generation lentiviral transfer plasmid encoding a GFP- or ALFA-tagged protein of interest and 2^nd^ generation packaging (e.g. psPAX2, Addgene #12260) and envelope plasmids (e.g. pMD2.G Addgene #12259). Transfected cells secrete lentiviral particles into the culture medium that are pseudotyped with the VSV-G envelope protein, which binds the widely expressed LDL receptor on the target cell surface and thus confers broad host cell tropism.

Lentiviral particles can then be used to transduce multiple human suspension and adherent cell lines that are compatible with our purification protocol. We favor a very rapid lentiviral transduction approach^1^ to generate Expi293F cell lines expressing a GFP or ALFA-tagged protein of interest. This approach is fast, easy to use and generates stable cell lines that robustly express a protein of interest. We prefer Expi293F (TetR-) and Expi293F inducible cell lines (TetR+) (Thermo Fisher Scientific, USA), because they maintain high viability even at very high cell density and thus maximize the yield of cells per liter of culture medium. To express secreted or membrane proteins without complex N-glycans, the N-acetylglucosaminyltransferase I knockout (GnTI-) derivatives of these cell lines should be used.

During infection, lentiviral particle-containing culture medium is simply mixed with human Expi293F suspension cells. This leads to the stable integration of the open-reading frame encoded on the transfer plasmid into a random genomic location. For first small-scale purification trials, we recommend expanding transduced cells to medium density in ∼50 ml. Once optimal conditions are found, purifications from ∼1-2 L high density cultures often provide enough material for structural and functional assays.

If lentiviral work is not feasible, multiple alternative approaches can be used to instead generate human suspension cell lines expressing a GFP- or ALFA-tagged protein of interest. These include for example transient transfection using polyethylenimine or Expifectamine (Thermo Fisher Scientific, USA), baculovirus transduction^15^ and adaptation of stable adherent cell lines to suspension growth^16^. Such stable adherent cell lines can be generated by Flp-In recombination^17^, antibiotic selection^16^ or CRISPR knock-in^18–20^.

### Cell lysis method and purification conditions

The subcellular localization and stability of the protein of interest dictates the appropriate lysis method to prepare cell extracts for affinity purification. Mechanical lysis is appropriate to purify soluble cytosolic or nuclear proteins. Nuclear DNA-bound proteins additionally need to be dissociated from DNA using a high salt lysis buffer^21^. Membrane proteins must be solubilized from whole cells or enriched membrane fractions using detergents or other membrane mimetics^22, 23^. The choice of detergent for solubilization can affect membrane protein extraction efficiency and stability.

Another critical factor to consider is the protein’s intrinsic stability. In general, lysis buffers with slightly elevated salt concentration (> ∼150 mM) and detergent can strongly reduce non-specific binding of lysate proteins to affinity resins. For example, the widespread RIPA lysis buffer contains high concentrations of ionic (deoxycholate and SDS) and non-ionic detergents (Triton-X-100 or NP-40) and is used for both soluble and membrane protein purification. However, some proteins and protein complexes can be very sensitive to high ionic strength or detergents. To determine the optimal conditions for a protein or complex of interest, we recommend optimizing lysis buffer composition in pilot small-scale purification trials.

### Protein purification by nanobody capture and protease release

The major components of our purification strategy are highly stable proteins that are compatible with most commonly used lysis buffer compositions.

For example, we use the anti-GFP nanobody Enhancer (12.8 kDa), which binds GFP with high affinity (590 pM) and specificity in many different systems^24, 25^. Various widespread GFP variants (including wildtype GFP, EGFP and superfolder GFP^26^), as well as closely related fluorescent proteins are compatible (**Extended Data Figure 3**). The ALFA nanobody (13.4 kDa) binds a rationally designed, 15 amino acid α-helical peptide with very high affinity (26 pM)^3^. Both nanobodies, as well as GFP and ALFA tags are stable and bind efficiently even under harsh conditions, including high concentrations of most detergents, salt and even urea (**Extended Data Figure 4**).

For our purification strategies, both nanobodies are immobilized onto Streptavidin beads using a fused biotin acceptor peptide (Avi tag), which is modified with a single biotin by the biotin ligase BirA either during expression in *E. coli* or after purification *in vitro*^27–29^. As one of the strongest non-covalent interactions in nature, the biotin-streptavidin interaction withstands harsh purification conditions.

Finally, our protocol makes use of the SUMO^Eu^ protein (10.5 kDa), which was engineered to be resistant against cleavage by endogenous eukaryotic SUMO proteases^4^. SUMO^Eu^-tagged nanobodies or proteins are thus stable in a variety of eukaryotic cells and cell extracts. The more commonly used yeast SUMO is useful for expression in prokaryotic systems, but is very rapidly cleaved in eukaryotic cells and cell lysate^4, 30^. The engineered SUMO^Eu^•SENP^EuB^ pair therefore extends the use of the SUMO protease technology to eukaryotes. A similar, but completely orthogonal SUMOstar system^31^ is commercially available (LifeSensors, USA), but slightly less resistant against host SUMO protease cleavage^4^.

### Preparation of the nanobody fusion proteins and SENP^EuB^ protease

All recombinant proteins required for this protocol can easily be purified in very high yield and purity from single 1 L *E. coli* cultures. Such preparations generate sufficient reagents for hundreds of purifications from eukaryotic cells. We provide expression plasmids and protocols for the generation of SENP^EuB^ protease, as well as biotinylated anti-GFP and anti-ALFA nanobodies with or without the SUMO^Eu^ module in between the Avi-tag and nanobody (see Table 1 for Addgene IDs).

The anti-GFP nanobody can easily be biotinylated during expression in the *E. coli* strain CVB101 (Avidity LLC, USA), which allows IPTG-inducible co-expression of the biotin ligase BirA. The anti-ALFA nanobody, however, expresses better in the *E. coli* Rosetta-gami2 strain (Millipore-Sigma, USA), which does not express BirA. Purified ALFA nanobody therefore needs to be biotinylated with purified BirA in solution. For this, we provide a BirA expression plasmid (pTP264) and purification protocol. BirA can, however, also be obtained commercially (Avidity LLC, USA).

Affinity resins containing immobilized (non-cleavable) anti-GFP or ALFA nanobodies are commercially available (ChromoTek, Germany; NanoTag Biotechnologies, Germany) and are directly compatible with the TagOFF strategy.

### Controls

If our protocol is to be used for the identification of interaction partners via mass spectrometry, it is important to include controls to account for trace contaminants that non-specifically bind to the affinity tag, nanobody or affinity resin. A good control is a matched cell lysate expressing solely the affinity tag without a fused protein i.e. only GFP (e.g. generated using plasmid pTP924). Less optimally, the cell lysate containing the protein of interest can be split into two even fractions and additionally incubated with beads that either do not contain any immobilized nanobody or that contain a non-specific nanobody, e.g. anti-ALFA nanobody for a GFP-tagged protein.

Similarly, for functional assays, it is important to control that the observed activity of the purified protein is not caused by a contaminant. If possible, a catalytically dead mutant should be purified in parallel under identical conditions.

The sorting of cell lines transduced with multiple different lentiviruses, encoding compatible fluorescent proteins, might require multicolor compensation to correct for cross-channel fluorescence bleed-through. Non-transduced, as well as single color control cells are typically required for compensation and also allow efficient sort gate placement. Plasmids pTP341 and pTP924, which express TagBFP and GFP, respectively, from a constitutive CMV promoter, can be used for this purpose.

### Limitations

One limitation of the lentiviral transduction approach is that the ectopic expression of a protein complex subunit can in some cases result in its purification in excess over other endogenous complex members. This especially affects subunits that are stable in the unassembled state. Frequently, these excess subunits can be separated by an additional size-exclusion chromatography purification step. However, multiple preventative measures can improve protein complex yield and stoichiometry. 1) Freestyle 293-F suspension cells can be used instead of Expi293F cells (Thermo Fisher Scientific, USA). We observed that overexpression of unassembled subunits is more limited in these cells, likely due to tighter regulation by the ubiquitin-proteasome system. However, Freestyle 293-F cells cannot be grown to as high density as Expi293F cells, decreasing yield per liter of expression. 2) Because our lentiviral vectors are regulated by doxycycline-inducible promoters, the expression level can also be fine-tuned by varying induction time in Expi293F TetR+ cells. 3) The expression level can be reduced by exchanging the strong CMV promoter for a weaker one^32^ and/or by removing the expression-enhancing WPRE element. 4) For membrane proteins it can be beneficial to first enrich a membrane fraction to remove excess non-incorporated and aggregated subunits in the cytosol. Purification from the solubilized membrane fraction can therefore yield higher sample quality and better complex stoichiometry.

If these approaches do not yield the expected outcome, it is possible to overexpress multiple subunits in parallel. This can be achieved in three different ways: 1) via co-transduction with multiple lentiviral particles, 2) via fusion of multiple subunits into a single expression plasmid using P2A sites or 3) via a combination of 1-2. The efficiency of dual or triple transduction is typically lower than transduction with a single lentivirus. Multi-color sorting via FACS is therefore required to obtain a fully transduced cell line. Our lentiviral transfer plasmid toolbox already allows generating compatible GFP and BFP expression plasmids. A third compatible plasmid can easily be generated by introducing mCherry.

Constitutive ectopic expression of a protein can be toxic or reduce cellular fitness. In a heterogenous cell population containing untransduced cells, this can lead to a gradual loss of more slowly growing transduced cells. In such cases, using the inducible expression system is likely to alleviate the negative selection pressure resulting from the constitutive expression of a toxic protein. Alternatively, successfully transduced cells can be sorted via FACS to remove faster growing untransduced cells.

The yield of lentiviral particles decreases sharply with increasing transfer plasmid size, resulting in strongly reduced titers for transfer plasmids encoding large proteins e.g. inserts between 5’ and 3’ Long-terminal repeats (LTRs) above 8 kbp. After subtraction of essential elements this leaves around 5 kbp for the GFP- or ALFA-tagged protein of interest (∼180 kDa). If necessary, removal of the WPRE element can yield an extra ∼600 bp.

### Expertise needed to implement the protocol

Our purification protocol requires only basic skills in recombinant protein production in *E. coli*, as well as cell culture handling and can even be carried out by qualified undergraduate students with prior training. The generation of stable human suspension cell lines using lentiviral transduction, however, requires special biosafety training and conditions in compliance with the relevant institutional and governmental biosafety regulations. Elegheert and colleagues expertly summarized common safety practices associated with lentiviral work^1^. However, if use of lentivirus is not feasible, multiple other strategies for plasmid delivery can be used instead (see above).

## Materials

### Cell lines

- Lenti-X 293T cell line (Takara Bio Inc., Japan; cat. no. 632180)
- Gibco Expi293F cells (TetR-; Thermo Fisher Scientific, USA; cat. no. A14527)
- Gibco Expi293F GnTI- cells (TetR-; Thermo Fisher Scientific, USA; cat. no. A39240)
- Gibco Expi293F inducible cells (TetR+; Thermo Fisher Scientific, USA; cat. no. A39241)
- Gibco Expi293F inducible GnTI- cells (TetR+; Thermo Fisher Scientific, USA; cat. no. A39242)
- Gibco Freestyle 293-F cell line (Thermo Fisher Scientific, USA; cat. no. R79007)

### E. coli strains

- NEBExpress I^q^ chemically competent cells (New England Biolabs, USA; cat. no. C3037I)
- CVB101 chemically competent cells (Avidity, USA)
- Stellar competent cells (Takara Bio Inc., Japan; cat. no. 636763)

**CRITICAL** Cloning DNA with repetitive elements, like our lentiviral transfer plasmid toolbox, requires a recA- *E. coli* strain like *E. coli* Stellar.

- Rosetta-gami 2 competent cells (Novagen / Millipore-Sigma, USA; cat. no. 71350-3)

### Reagents

- Gibco Expi293 Expression (Expi) Medium (Thermo Fisher Scientific, USA; cat. no. A14351-01)
- Gibco FreeStyle 293 expression medium (Thermo Fisher Scientific, USA; cat. no. 12338018)
- Gibco DMEM, high glucose, no glutamine (Thermo Fisher Scientific, USA; cat. no. 11960051)
- Gibco L-glutamine 200 mM (100x Gln) (Thermo Fisher, cat. no. 25030081)
- Gibco Penicillin-Streptomycin 5000 U/mL (100x Pen-Strep) (Thermo Fisher Scientific, USA; cat. no. 15070063)
- HyClone fetal bovine serum (FBS) (Cytiva, USA; cat. no. SH30071.03)
- Gibco DPBS, no calcium, no magnesium (Thermo Fisher Scientific, USA; cat. no. 14190136)
- Gibco Trypsin-EDTA (0.25%), phenol red (Thermo Fisher Scientific, USA; cat. no. 25200056)
- Dimethyl Sulfoxide (DMSO) (Thermo Fisher Scientific, USA; cat. no. A13280.36)
- Gibco Opti-MEM I (1x) Reduced Serum Medium (Thermo Fisher Scientific, USA; cat. no. 31985-062)
- TransIT-293 Transfection Reagent (Mirus Bio, USA; cat. no. MIR 2704)
- LB Broth, Miller (Fisher Scientific, USA; cat. no. BP1426-2)
- LB Agar, Miller (Powder) (Fisher Scientific, USA; cat. no. BP1425-500)
- Tryptone (Fisher Scientific, USA; cat. no. BP9726-5)
- Yeast extract, granulated (Fisher Scientific, USA; cat. no. BP9727-5)
- SOC Recovery Medium (Thermo Fisher Scientific, USA; cat. no. 15544034)
- Carbenicillin (Disodium Salt) (Fisher Scientific, USA; cat. no. BP26485)
- Kanamycin Sulfate (Fisher Scientific, USA; cat. no. BP906-5)
- Chloramphenicol (Fisher Scientific, USA; cat. no. BP904-100)
- IPTG (Millipore-Sigma, USA; cat. no. I6758-5G)
- Imidazole (Millipore-Sigma, USA; cat. no. I2399-500G)
- NaCl (Fisher Scientific, USA; cat. no. S9888-5KG)
- Tris Base, Trizma (Millipore-Sigma, USA; cat. no. T6066-5KG)
- HEPES (Millipore-Sigma, USA; cat. no. H3375-500G)
- 1 M Magnesium Acetate solution (Millipore-Sigma, USA; cat. no. 63052-100ML)
- Magnesium Chloride (VWR, USA; cat. no. MK5958-04)
- Potassium Acetate (Millipore-Sigma, USA; cat. no. 60035-1KG)
- Hydrochloric acid, concentrated 37%/12N (Millipore-Sigma, USA; cat. no. HX0603-3)
- Potassium hydroxide pellets (VWR, USA; cat. no. 6984-06)
- Glycerol (Fisher Scientific, USA; cat. no. BP229-4)
- Sucrose (Millipore-Sigma, USA; cat. no. 50389-5KG)
- DTT (Millipore-Sigma, USA; cat. no. D9163-25G)
- Ethanol 200 Proof (VWR, USA; cat. no. TX-89125172CAL)
- Methanol (VWR, USA; cat. no. BDH1135-4LG)
- PMSF (Thermo Fisher Scientific, USA; cat. no. 36978)
- Ribonuclease A from bovine pancreas (Millipore-Sigma, USA; cat. no. R6513-50MG)
- Adenosine 5′-triphosphate dipotassium salt hydrate (Millipore-Sigma, USA; cat. no. A8937-1G)
- D-Biotin (Millipore-Sigma, USA; cat. no. B4501-1G)
- dPEG24-biotin acid (Quanta Biodesign, USA; cat. no. 10773)
- Triton-X-100 (Millipore-Sigma, USA; cat. no. X100-500)
- Tween-20 (Millipore-Sigma, USA; cat. no. P1379-1L)
- GDN (Anatrace, USA; cat. no. GDN101 25 GM)
- Doxycycline hyclate (Millipore-Sigma, USA; cat. no. D9891-1G)

• 20 % (w/v) SDS (VWR, USA; cat. no. 97062-442)

- Bromophenol Blue (Millipore-Sigma, USA; cat no. B0126-25G)
- Urea (Millipore-Sigma, USA; cat. no. U0631-500G)
- Roche cOmplete, Mini, EDTA-free protease inhibitor cocktail (Millipore Sigma, USA; cat. no. 11836170001)
- Ni-NTA agarose resin 25 ml (Qiagen, Germany; cat. no. 30210)
- Pierce Streptavidin magnetic beads (Thermo Fisher Scientific, USA; cat. no. 88816)

### Plasmids

- pMD2.G lentivirus envelope plasmid (Addgene ID #12259)
- psPAX2 lentivirus packaging plasmid (Addgene ID #12260)

### Sequencing primers

- CMV forward (5’– CGCAAATGGGCGGTAGGCGTG –3’)
- WPRE reverse (5’– GTTGCCTGACAACGGGCC –3’)
- GFP C-terminus forward (5’– GGAGACGGTCCCGTCCTC –3’)
- BFP C-terminus forward (5’– GATACTGCGACCTCCCTAGC –3’)
- GFP N-terminus reverse (5’– TGGCCATTCACGTCTCCGTC –3’)
- BFP N-terminus reverse (5’– CTTGAAGTGATGGTTGTCCACGGTGC –3’)

### Equipment

- 125 mL Erlenmeyer flask, vent cap, plain bottom, PETG (Celltreat, cat. no. 229801)
- 490 cm^2^ tissue culture treated roller bottle, vented cap (1 L roller bottle; Celltreat, cat. no. 229383)
- 850 cm^2^ tissue culture treated roller bottle, vented cap (2 L roller bottle; Celltreat, cat. no. 229385)
- Tissue culture 6-well plates (Genesee Scientific, USA; cat. no. 25-105)
- Tissue culture 150 mm plates (Fisher Scientific, USA; cat. no. FB012925)
- 5 ml serological (Genesee Scientific, USA; cat. no. 12-102)
- 10 ml serological (Genesee Scientific, USA; cat. no. 12-104)
- 25 ml serological (Genesee Scientific, USA; cat. no. 12-106)
- 50 ml serological (Genesee Scientific, USA; cat. no. 12-107)
- Steriflip-GP Sterile centrifuge tube top filter unit (Millipore Sigma, cat. no. SCGP00525)
- 500 ml sterile filter units (Genesee Scientific, USA; cat. no. 25-227)
- Corning cryogenic vials, external thread (2.0 mL; Millipore Sigma, cat. no. CLS430661)
- 5 ml microcentrifuge tube, sterile (Eppendorf, Germany; cat. no. 0030119487)
- 15 ml disposable centrifuge tube, sterile (Fisher Scientific, USA; cat. no. 05-539-12)
- 50 ml disposable centrifuge tube, sterile (Fisher Scientific, USA; cat. no. 05-539-8)
- FlowTubes with strainer cap (VWR, USA; cat. no. 76449-658)
- Tissue culture CO2 incubator (Thermo Fisher Scientific, USA; model no. Heracell 240i)
- Tissue culture S41i CO2 Incubator Shaker (Eppendorf, Germany; cat. no. S41I120010)
- Class II, type A2 biosafety cabinet (Baker, USA; model: SterilGard III SG403)
- Cell sorter (Sony Biotechnology, USA; model no. SH800S)
- Automated cell counter (Thermo Fisher Scientific, USA; model no. Countess 3)
- Cell counting slides (Bulldog-bio, USA; cat. no. DHC-N01)
- Tissue culture Floid cell imaging station (Thermo Fisher Scientific, USA; cat. no. 4471136)
- Benchtop centrifuge model no. 5810 with swing bucket rotor S-4-104 (Eppendorf, Germany; cat. no. 022627110) including 4x 750 ml swing buckets + 15 ml conical tube adapters, as well as matching aerosol-tight caps (cat. no. 022638661) for lentiviral harvest.
- Refrigerated floor centrifuge (Thermo Fisher Scientific, USA; model no. Sorvall RC6+) with Fiberlite F9-4x1000y rotor
- Corning CoolCell cell freezing vial container (Fisher Scientific, USA; cat. no. 07-210-002)
- Fast performance liquid chromatography system (Cytiva, USA; model no. Äkta Pure 25 M)
- Size exclusion chromatography column (Cytiva, USA; model no. Superose 6. Increase 3.2/300)
- Hamilton gas-tight syringe 50 µl (Millipore-Sigma, USA; cat. no. 26280-U)
- Ultra centrifugal filters for protein concentration (Millipore-Sigma, USA; model no. Amicon Ultra 0.5 or 4 with protein-specific molecular weight cut-off)
- Disposable chromatography column, 20 ml (Bio-Rad, USA; cat. no. 7321010)
- PD-10 desalting columns (Cytiva, USA; cat. no. 17085101)
- Wheaton Dounce tissue grinder, 15 mL, tight pestle (DWK Life Sciences, USA; cat. no. 357544)
- Branson Ultrasonics sonifier SFX250 cell disruptor (Fisher Scientific, USA; cat. no. 15-345-138) with disruptor horn (cat. no. 22-020860)
- Invitrogen DynaMag-2 Magnet (Thermo Fisher Scientific, USA; cat. no. 12321D)
- Invitrogen DynaMag-15 Magnet (Thermo Fisher Scientific, USA; cat. no. 12301D) (cost-efficient alternatives are available here: https://sergilabsupplies.com)

### Reagent setup

**LB Broth (LB):** To make 1 L of LB weigh in 25 g of LB Broth and dissolve in 1 L ddH2O. Sterilize by autoclaving for 45 min. at 121°C using liquid cycle.

**Super Broth (SB):** To prepare 5 L of SB weigh in 175 g tryptone, 100 g yeast extract, 25 g NaCl and dissolve in 4.5 L ddH2O. Adjust pH to 7-7.5 with 1M NaOH and top up to 5 L. Aliquot 1 L into 2 L Erlenmeyer flasks and sterilize by autoclaving for 45 min. at 121°C using liquid cycle.

**LB agar plates:** Dissolve 25 g LB Agar in 1 L ddH2O. Sterilize by autoclaving for 45 min. at 121°C using liquid cycle. When cooled to 45-50°C add 1 ml of a 1000x stock of antibiotic, mix, and pour into 100x15 mm petri dishes. Allow to solidify and dry at room temperature.

**50 mg/ml Kanamycin (Kan) (1000x stock):** To prepare 10 ml of a 50 mg/mL stock solution of kanamycin, dissolve 0.5 g of kanamycin sulfate in 9.5 mL of H2O. Top up the volume to 10 mL and filter-sterilize using a 0.22-μm filter. Store in 1 mL aliquots at −20°C.

**100 mg/ml Carbenicillin (Carb) (1000x stock):** To prepare 10 ml of a 100 mg/mL stock solution of carbenicillin, dissolve 1 g of carbenicillin (disodium salt) in 9.5 mL of H2O. Top up the volume to 10 mL and filter-sterilize using a 0.22-μm filter. Store in 1 mL aliquots at −20°C.

**50 mg/ml Chloramphenicol (Cam):** To prepare 10 ml of a 50 mg/mL stock solution of chloramphenicol, dissolve 0.5 g of chloramphenicol in 9.5 mL 100 % ethanol. Top up the volume to 10 mL and filter-sterilize using a 0.22-μm filter. Store in 1 mL aliquots at −20°C.

- **M IPTG:** To make up 10 ml of 1 M IPTG, weigh in 2.38 g of IPTG and dissolve in 8 ml ddH2O. Top up the volume to 10 mL and filter-sterilize using a 0.22-μm filter. Store in 1 mL aliquots at −20°C.
- **M Tris/HCL pH 7.5:** To make 1 L of 2 M Tris/HCL at pH 7.5, weigh in 242.2 g Tris base and dissolve in 800 ml ddH2O. Add 134.3 ml concentrated 37%/12N HCL. Top up the volume to 1 L with ddH2O and filter-sterilize. Store at RT.

**1 M HEPES/KOH pH 7.5:** To make 0.5 L of 1 M HEPES/KOH at pH 7.5, weigh in 119.2 g HEPES and dissolve in 350 ml ddH2O. Adjust pH to 7.5 with potassium hydroxide pellets. Top up the volume to 0.5 L with ddH2O and filter-sterilize. Store at 4°C.

**5 M NaCl:** To make 1 L of 5 M NaCl, dissolve 292 g of NaCl in 700 ml ddH2O and add ddH2O up to 1 L. Filter-sterilize and store at RT.

**5 M KAcetate (KAc):** To make 200 ml of 5 M KAc, dissolve 98.14 g of KAc in 180 ml ddH2O and add 800 µl concentrated 37%/12 N HCl. Add ddH2O up to 200 ml and filter-sterilize. Store at RT.

- **M MgCl2:** To make 100 ml of 1 M MgCl2, dissolve 20.3 g of MgCl2 in 80 ml ddH2O and add ddH2O up to 100 ml. Filter-sterilize and store at RT.
- **M Imidazole pH 7.5:** To make 250 ml of 2 M imidazole pH 7.5, dissolve 34.04 g imidazole in 200 ml ddH2O. Adjust pH to 7.5 with concentrated 37%/12N HCl and add ddH2O up to 250 ml. Filter-sterilize and store at 4 C°, protected from light.

**1 M DTT:** To make 10 ml of 1 M DTT, dissolve 1.5 g of DTT in 8 ml ddH2O and add ddH2O up to 10 ml. Make 1 ml aliquots and store at -20°C.

**100 mM PMSF:** To make 50 ml of 100 mM PMSF, weigh in 0.87 g PMSF and dissolve in 50 ml methanol. Store at - 20°C. **! CAUTION** Both PMSF and methanol are toxic.

**100 mM ATP:** To make 10 ml of 100 mM ATP, weigh in 551 mg of Adenosine 5′-triphosphate disodium salt hydrate and dissolve in 8 mL ddH2O. Adjust to pH 7.0 with potassium hydroxide pellets. Top up the volume to 10 mL with ddH2O and make 0.5 ml aliquots. Store at -20°C.

**10 mg/ml RNase A:** To make 5 ml of a 10 mg/ml stock of RNaseA, reconstitute 50 mg lyophilized RNase A in 5 ml 1x PBS and freeze in 1 ml aliquots, store at -20°C

**50 mM biotin stock:** To make 10 ml of a 50 mM biotin stock in 50 mM HEPES/KOH pH 7.5, weigh in 122 mg biotin and dissolve in 10 ml 50 mM HEPES/KOH pH 7.5. Make 1 ml aliquots and store at -20°C.

**50 mM dPEG24-biotin acid stock:** To make 1 ml of a 50 mM dPEG24-biotin acid stock in 50 mM HEPES/KOH pH 7.5, weigh in 68.6 mg dPEG24-biotin acid and dissolve in 1 ml 50 mM HEPES/KOH pH 7.5. Make 100 µl aliquots and store at -20°C.

**8 M Urea:** To prepare 50 ml of 8 M Urea, weigh in 24 g Urea and dissolve in 20 ml ddH2O. Add ddH2O up to 50 ml, filter-sterilize and store at RT.

**1 mg/ml doxycycline**: To prepare a 50 ml of 1 mg/ml doxycycline, weigh in 50 mg doxycycline and dissolve in 50 ml ddH2O. Filter-sterilize using a 0.22-μm filter in a tissue culture hood. Make 1 ml aliquots and store at -20°C. For the induction of larger culture volumes a separate 10 mg/ml stock is useful.

**25x Protease inhibitor cocktail:** Dissolve 1x Roche cOmplete, Mini, EDTA-free protease inhibitor cocktail tablet in 1 ml ddH2O. Use immediately or store at -20°C.

**10 % (w/v) GDN:** To prepare 10 ml 10 % (w/v) GDN, weigh in 1 g GDN and add 8 ml ddH2O. Rotate head-over-tail at RT until GDN is fully dissolved. Top up to 10 ml with ddH2O and make 1 ml aliquots. Store at -20°C.

**20 % (v/v) Triton-X-100:** To prepare 100 ml of 20 % (v/v) Triton-X-100, transfer 20 ml 100 % (v/v) Triton-X-100 into a 100 ml glass bottle and add 80 ml ddH2O. Incubate tumbling or rotating until the solution is well mixed. Store at RT.

**5x SDS-PAGE sample buffer:** 250 mM Tris/HCl pH 6.8, 5 % (w/v) SDS, 50 % (v/v) glycerol, 500 mM DTT. Prepare 40 ml by mixing the following: 1.211 g Tris base, 10 ml 20% (w/v) SDS, 20 ml 100 % (v/v) glycerol and 3.084 g DTT. Adjust pH to 6.8 by adding 0.79 ml 37%/12N HCl and then add ddH2O to 40 ml. Mix by inverting gently until all DTT has dissolved and then add 40 mg Bromophenol Blue. Mix as above and make 1 ml aliquots. Store at -20°C.

**Resuspension buffer:** 50 mM Tris/HCl pH 7.5, 300 mM NaCl, 20 mM imidazole, 1 mM DTT, 1 mM PMSF. Prepare 1 L by mixing the following: 25 ml 2 M Tris/HCl pH 7.5, 60 ml 5 M NaCl, 10 ml 2 M imidazole and 905 ml ddH2O. Store at 4°C. Add 1 mM DTT and 1 mM PMSF directly before use.

**Low-salt-biotin-ATP-RNase A (LSBAR) buffer**: 50 mM HEPES/KOH pH 7.5, 100 mM KAc, 2 mM MgAc, 1 mM DTT, 1 mM ATP, 50 µM biotin and 100 µg/ml RNAse A. Prepare 50 ml by mixing the following: 2.5 ml HEPES/KOH pH 7.5, 1 ml 5 M KAc, 0.5 ml 100 mM ATP, 50 µl 50 mM biotin, 0.5 ml 10 mg/ml RNase A, 50 µl 1 M DTT and 45.4 ml ddH2O. Prepare fresh shortly before use.

**Low-salt-ATP (LSA) buffer**: 50 mM HEPES/KOH pH 7.5, 100 mM KAc, 2 mM MgAc, 1 mM DTT, 1 mM ATP. Prepare 50 ml by mixing the following: 2.5 ml HEPES/KOH pH 7.5, 1 ml 5 M KAc, 0.5 ml 100 mM ATP, 50 µl 1 M DTT and 45.95 ml ddH2O. Prepare fresh shortly before use.

**High-salt buffer:** 50 mM Tris/HCl pH 7.5, 1 M NaCl, 20 mM imidazole, 1 mM DTT. Prepare 50 ml by mixing the following: 1.25 ml 2 M Tris/HCl pH 7.5, 10 ml 5 M NaCl, 0.5 ml 2 M imidazole and 38.2 ml ddH2O. Store at 4°C. Add 50 µl 1 M DTT directly before use.

**Imidazole elution buffer:** 50 mM Tris/HCl pH 7.5, 300 mM NaCl, 500 mM imidazole, 10% glycerol, 1 mM DTT. Prepare 50 ml by mixing the following: 1.25 ml Tris/HCl pH 7.5, 3 ml 5 M NaCl, 12.5 ml 2 M imidazole pH 7.5, 5 ml 100 % glycerol and 28.2 ml ddH2O. Store at 4°C. Add 50 µl 1 M DTT directly before use.

**BirA storage buffer**: 50 mM Tris/HCl pH 7.5, 200 mM NaCl, 1 mM DTT, 250 mM sucrose. Prepare 50 ml by mixing the following: 1.25 ml Tris/HCl pH 7.5, 2 ml 5 M NaCl, 6.58 ml 1.89 M/65% (w/v) sucrose and 40.12 ml ddH2O. Store at 4°C. Add 50 µl 1 M DTT directly before use.

**ALFA Nb storage buffer:** 50 mM Tris/HCl pH 7.5, 300 mM NaCl, 250 mM sucrose. Prepare 50 ml by mixing the following: 1.25 ml Tris/HCl pH 7.5, 3 ml 5 M NaCl, 6.58 ml 1.89 M/65% (w/v) sucrose and 39.17 ml ddH2O. Store at 4°C.

**5x biotinylation buffer:** 250 mM HEPES/KOH pH 7.5, 500 mM NaCl, 50 mM ATP, 62,5 mM MgCl2 and 50 mM biotin. Prepare 10 ml by mixing the following: 1.25 ml 2 M Tris/HCl pH 7.5, 1 ml 5 M NaCl, 5 ml 100 mM ATP, 0.625 ml 1 M MgCl2 and 2.125 ml ddH2O. Weigh in 122 mg biotin and dissolve in 10 ml buffer as prepared above. Make 1 ml aliquots and store at -20°C.

**SA test binding (STB) buffer:** 50 mM Tris/HCl pH 7.5, 200 mM NaCl, 0.1 % (v/v) Triton-X-100, 1 mM DTT. Prepare 20 ml by mixing the following: 0.5 ml Tris/HCl pH 7.5, 0.8 ml 5 M NaCl, 0.1 ml 20 % (v/v) Triton-X-100, 20 µl 1 M DTT and 18.58 ml ddH2O. Prepare freshly before use.

**Solubilization buffer:** 50 mM HEPES/KOH pH 7.5, 200 mM NaCl, 2 mM MgAc, 1 mM ATP, 1 % (w/v) GDN, 1x Roche cOmplete protease-inhibitor cocktail, 1 mM DTT. Prepare 10 ml by mixing the following: 0.5 ml 1 M HEPES/KOH pH 7.5, 0.4 ml 5 M NaCl, 20 µl 1 M MgAc, 0.1 ml 100 mM ATP, 1 ml 10% (w/v) GDN, 0.4 ml 25x Roche cOmplete protease-inhibitor cocktail, 10 µl 1 M DTT and 7.57 ml ddH2O. Prepare freshly before use.

**Wash buffer:** 50 mM HEPES/KOH pH 7.5, 200 mM NaCl, 2 mM MgAc, 0.5 mM ATP, 0.01 % (w/v) GDN, 1x Roche cOmplete protease-inhibitor cocktail, 1 mM DTT. Prepare 10 ml by mixing the following: 0.5 ml 1 M HEPES/KOH pH 7.5, 0.4 ml 5 M NaCl, 20 µl 1 M MgAc, 0.05 ml 100 mM ATP, 10 µl 10% (w/v) GDN, 0.4 ml 25x Roche cOmplete protease-inhibitor cocktail, 10 µl 1 M DTT and 8.61 ml ddH2O. Prepare freshly before use.

**Wash buffer (-ATP):** 50 mM HEPES/KOH pH 7.5, 200 mM NaCl, 2 mM MgAc, 0.01 % (w/v) GDN, 1x Roche cOmplete protease-inhibitor cocktail, 1 mM DTT. Prepare 1 ml by mixing the following: 0.05 ml 1 M HEPES/KOH pH 7.5, 0.04 ml 5 M NaCl, 2 µl 1 M MgAc, 1 µl 10% (w/v) GDN, 0.04 ml 25x Roche cOmplete protease-inhibitor cocktail, 1 µl 1 M DTT and 0.866 ml ddH2O. Prepare freshly before use.

**DMEM/10%FBS/1xGln medium (DMEM):** To 500 mL Gibco DMEM, high glucose, no glutamine add 50 mL FBS and 5.5 mL 100x Gln. (Optional) Add 5.5 mL 100x Pen-Strep.

### Procedure

#### Generation of biotinylated nanobodies and SENP^EuB^

##### Preparation of biotinylated GFP nanobodies ● Timing ∼4d

Steps 1-12 describe the expression of biotinylated anti-GFP nanobodies from plasmids pTP396 and pTS117 in *E. coli* CVB101. (Optional) Take samples for SDS-PAGE to help troubleshoot protein expression and purification as outlined in **Supplementary Data 2**.

**CRITICAL STEP** The ALFA nanobody does not express well in *E. coli* CVB101 and thus needs to be expressed in an alternative strain and biotinylated in vitro (see below).

1 **Transformation via heat-shock**: Thaw one vial of *E. coli* CVB101 cells on ice for 10 min. and then add 1 µl of ∼100 ng/µl plasmid. Mix and incubate for 30 min on ice. Heat-shock vial at 42°C for 30 sec and then quickly remove and incubate for 1 min on ice. Resuspend heat-shocked cells in 300 µl SOC recovery medium and shake at 37°C for 1 h. Plate 50 µl of transformed cells onto an LB agar plate containing 50 µg/ml Kan and 10 µg/ml Cam using the dilution streak method. Incubate plate at 37°C overnight.

**CRITICAL STEP** Chloramphenicol is required to maintain the BirA expression plasmid.

2 **Starting a pre-culture**: Pick a single colony from the plate and transfer into a 1.5 ml Eppendorf tube containing 200 µl SB supplemented with 50 µg/ml Kan and 10 µg/ml Cam (SB-Kan-Cam). Incubate shaking at 37°C for 4-5 h and then transfer to 100 ml SB-Kan-Cam in a 2-5 L baffled flask. Incubate this pre-culture at 37°C overnight with shaking at ∼220 rpm.

3 **Inducing the main culture**: The next morning, measure the optical density of a 1:10 dilution of the pre-culture at 600 nm (OD600) using a spectrophotometer. Dilute the pre-culture with SB-Kan-Cam to an OD600 of ∼1 to make up a 1 L main culture. Weigh in ∼12.2 mg biotin from powder, resuspend in 1 ml SB-Kan and add to main culture (∼50 µM final). Incubate main culture at 18°C with shaking for around 1 h. Induce main culture by addition of 0.2 mM IPTG and incubate shaking overnight (∼20 h) at 18°C.

**CRITICAL STEP** Biotin supplementation is crucial for efficient biotinylation of the expressed nanobodies.

4 **Harvest main culture**: Harvest cells by centrifugation at 9,220 g for 10 min. at 4°C. Discard supernatant and resuspend cell pellet in 120 ml resuspension buffer. Aliquot 4x 30 ml into 50 ml tubes and freeze in liquid nitrogen.

**! CAUTION** Wear safety goggles when handling liquid nitrogen.

**PAUSE POINT** Frozen *E. coli* cell pellets can be stored at -80°C for many months.

5 **Cell lysis**: Rapidly thaw cell pellets in lukewarm water and place on ice as soon as the last frozen clumps are thawed. Place cells into a ∼250 ml thin-walled metal beaker and transfer into an ice-water bath. Lyse cells by sonication on a Branson sonifier using the flat tip and 4x 1 min. sonication cycles at 100% amplitude. Each cycle should have roughly 1 sec pulses followed by 2 sec breaks. In between the four cycles mix lysate and allow to cool down for 30 sec - 1 min.

**CRITICAL STEP** Sonication creates heat that can activate *E. coli* proteases and denature proteins, causing degradation, aggregation or excessive chaperone binding. Keep thawed cells in an ice-water bath at all times. Ideally perform sonication in a 4°C cold room. Keep sonication probe properly submerged in the cell lysate to prevent foam formation (aggregated protein).

6 **Centrifugation:** Centrifuge the lysate for 30 min. at 35,000 g and 4°C. Take off and pool supernatant.

**PAUSE POINT** Lysate can be supplemented with either 250 mM sucrose or 10% glycerol and frozen in liquid nitrogen for storage at -80°C for many months depending on protein stability.

7 **Binding to Ni^2+^-resin:** Equilibrate around 2 ml settled Ni-NTA agarose resin with 20 ml resuspension buffer in a disposable 20 ml gravity flow column. Drip speed can be enhanced by attaching a long and wide, ideally blunt-ended needle to the column. Transfer equilibrated beads to a 100 ml glass bottle containing ∼60 ml lysate and incubate for 1 h at 4°C with constant mixing. We suggest either purifying 2x 60 ml lysate in two separate columns or just freezing 60 ml lysate for future purification.

8 **Washing the Ni^2+^-resin:** Transfer the suspension back to the column and discard the flow- through. Wash the resin with 20 ml of the following buffers in this order: 1) resuspension buffer, 2) LSBAR buffer, 3) high-salt buffer and finally 4) resuspension buffer.

9 **Elution:** Detach needle if used, apply 1 ml imidazole elution buffer and discard flow-through. Elute stepwise by adding 0.5 ml elution buffer to the resin. Collect a total of 6x 0.5 ml fractions in separate 1.5 ml tubes.

10 **Determine concentration:** Measure UV absorbance of each fraction at 280 nm and pool fractions with the highest protein content. Estimate the concentration of the pooled purified nanobody using its specific extinction coefficient at 280 nm (ε280) (see **Table 2**). Make sure to use elution buffer as a blank control. A typical yield for pTP396 is 60 mg per 1 L of bacterial culture. pTP396 can be stored in highly concentrated form, e.g. ∼600 µM stocks.

**Table 2.**
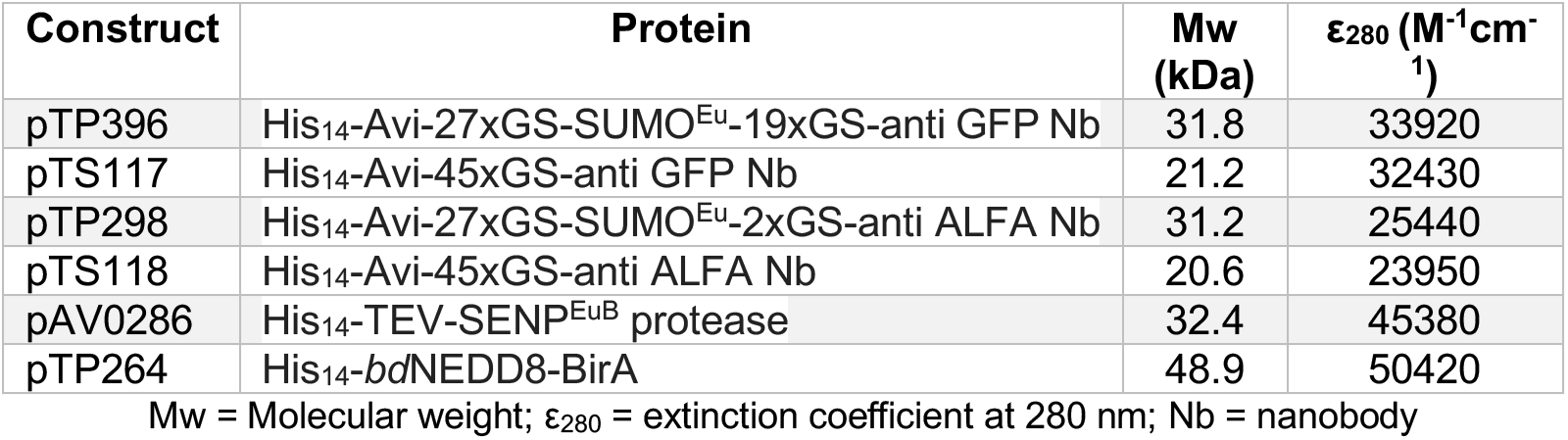
Properties of all purified proteins.

11 **SDS-PAGE**: Analyze expression and purification using SDS-PAGE and Coomassie staining. Compare purified protein to **Fig. 5a**.

**Fig. 5.**
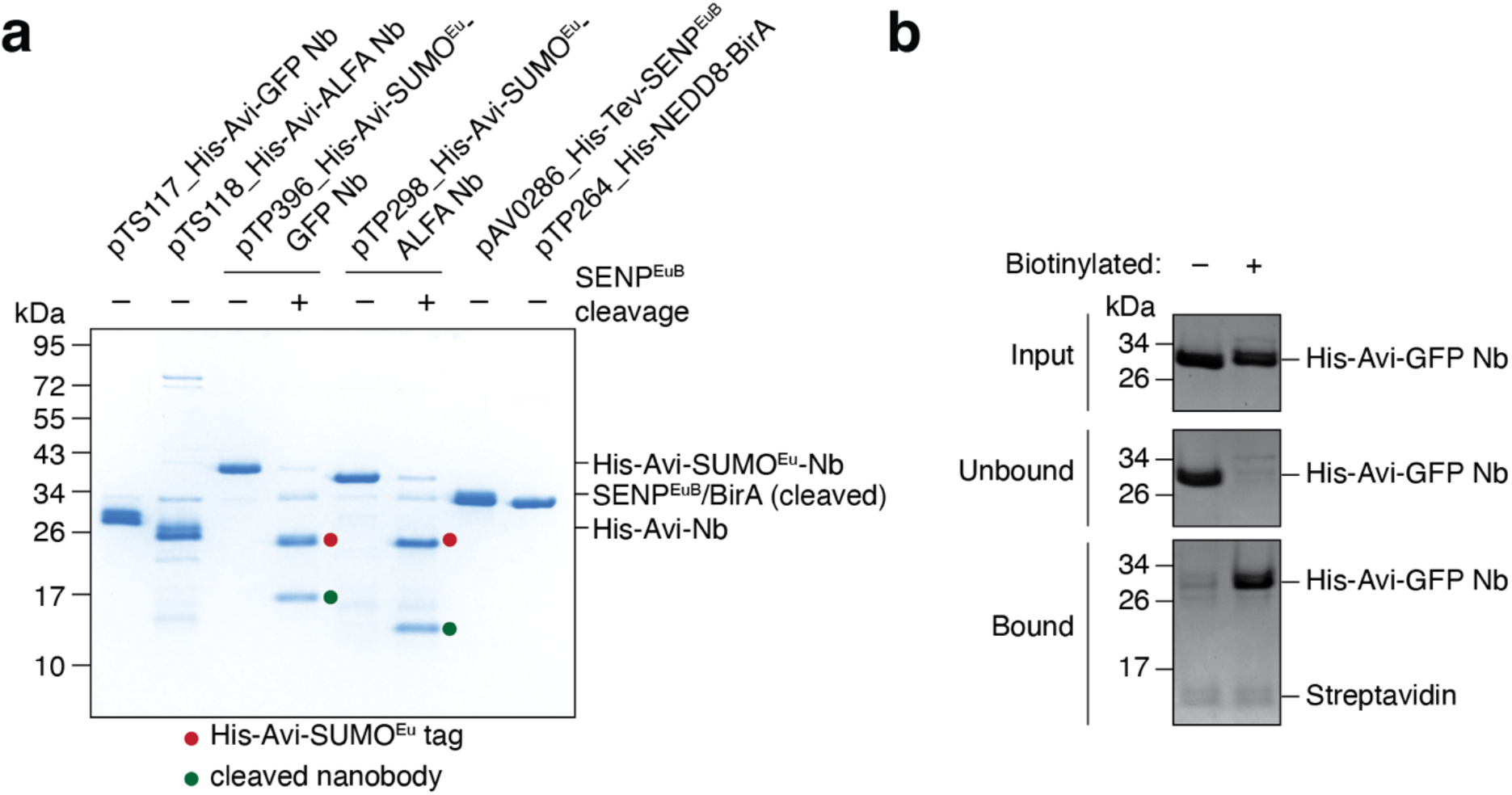
Quality control of purified proteins. (**a**) 1 µg of each of the protein reagents required for the protocol was analyzed by SDS-PAGE and Coomassie staining. pTP396 and pTP298 were additionally cleaved with SENP^EuB^ in solution to assess its activity. His-NEDD8-tagged BirA was purified by Ni^2+^-chelate affinity purification and eluted by cleavage with NEDP1 protease^33, 34^, resulting in untagged BirA. The removal of the His-NEDD8 tag is not required. Nb = nanobody (**b**) Coomassie-stained SDS-PAGE gel showing the quantitative streptavidin test binding of purified, biotinylated His-Avi-anti-GFP Nb (pTS117; steps 17-23). The protein was expressed and purified from *E. coli* NEBExpress (-biotinylation) or *E. coli* CVB101 (+biotinylation).

### ? TROUBLESHOOTING

12 **Freeze**: Aliquot purified nanobodies into multiple 10 and 50 µl aliquots in thin-walled 200 µl PCR tubes and flash freeze in liquid nitrogen. Store in small boxes or 50 ml tubes at -80°C. **PAUSE POINT** Frozen protein aliquots are stable at -80°C for years.

**CRITICAL STEP** Although aliquots typically tolerate multiple freeze-thaw cycles, repeated freeze-thawing or prolonged incubation at elevated temperatures will result in nanobody aggregation. Mark freeze-thaw cycles on the tubes and keep aliquots on ice immediately after thawing.

#### Preparation of SENP^EuB^ protease and biotin ligase BirA ● Timing ∼4 d

SENP^EuB^ and BirA are expressed from pAV0286 and pTP264, respectively, by following steps 1- 12 with the following modifications:

- Step 1: Use *E. coli* strain NEBExpress instead, as no biotinylation is required. Leave out Cam from the LB-Kan agar plate, pre-culture and main culture.
- Step 3: Induce protein expression for 6 hours at 18°C. There is no need to supplement the medium with biotin.
- Step 7: Use 2 ml Ni-NTA agarose beads and incubate with 120 ml lysate.
- Step 8: Replace LSABR buffer with LSA buffer (without biotin and RNase A).
- Steps 11-12: For SENP^EuB^ follow steps 11-12, since it can be stored in elution buffer. For BirA it is necessary to exchange the elution buffer to BirA storage buffer using a PD-10 desalting column. This is necessary because BirA activity is sensitive to ionic strength and glycerol concentration. Closely follow the manufacturer’s instructions. Pool peak fractions based on absorbance at 280 nm, measure the final concentration of the pooled fractions and analyze an aliquot by SDS-PAGE. Flash freeze aliquots of the stock in liquid nitrogen as described above.

#### Preparation of ALFA nanobodies ● Timing ∼4 d

ALFA nanobodies are expressed from pTP298 and pTS118 by following steps 1-12 with the following modifications:

- Step 1: Use *E. coli* strain Rosetta-gami 2 instead and supplement LB-Kan agar plates, pre- and main cultures additionally with 34 µg/ml Cam to maintain the pRARE2 plasmid. **CRITICAL STEP** This strain does not support biotinylation during expression. The resulting anti-ALFA tag nanobodies therefore need to be biotinylated with purified biotin ligase BirA *in vitro* after Ni^2+^-chelate affinity purification and buffer exchange as described below.
- Step 3: Induce protein expression for 6 hours at 18°C. There is no need to supplement the medium with biotin.
- Step 7: Use 2 ml Ni-NTA agarose beads and incubate with 120 ml lysate.
- Steps 11-12: Exchange the imidazole elution buffer after Ni^2+^-purification to ALFA Nb storage buffer using a PD-10 desalting column. Closely follow the manufacturer’s instructions, pool peak fractions based on absorbance at 280 nm, measure the final concentration and 260/280 nm absorbance ratio of the pooled fractions.

**PAUSE POINT** Either proceed straight to *in vitro* biotinylation or flash freeze 1 ml aliquots in liquid nitrogen.

#### *In vitro* biotinylation of ALFA nanobodies with purified BirA ● Timing ∼5 h

Ni^2+^-purified and buffer exchanged ALFA nanobodies expressed from pTP298 and pTS118 are biotinylated with BirA in solution. Excess biotin then needs to be removed by buffer exchange.

13 **Set up reaction:** Mix 100 nmoles of pTP298 or pTS118 with 1.5 nmoles of BirA enzyme, 300 µl 5x biotinylation buffer and add ddH2O to a final reaction volume of 1.5 ml. **CRITICAL STEP** *In vitro* biotinylation is most efficient when the substrate is provided at a concentration of at least 40 µM. Reaction volumes can be scaled up or down.

14 **Incubation:** Incubate *in vitro* biotinylation reaction for 3 h at 25°C in a thermomixer with only occasional mixing every 30 minutes.

**CRITICAL STEP** BirA activity is highly temperature-dependent. Biotinylation reactions on ice typically need at least 24 h to reach completion.

15 **Removal of excess biotin:** Load the entire biotinylation reaction on a PD-10 desalting column, pre-equilibrated in ALFA nanobody storage buffer according to the manufacturer’s instructions. Let all 1.5 ml completely sink into the column and discard the flow-through. Then add 1 ml of ALFA nanobody storage buffer and discard the flow-through. Now elute stepwise by adding 0.5 ml ALFA nanobody storage buffer to the PD-10 desalting column and collect 6x 0.5 ml fractions in separate 1.5 ml tubes.

16 **Pool biotinylated nanobody fractions:** Measure the absorbance at 280 nm, as well as the 260/280 nm absorbance ratio of the individual fractions. Pool only fractions with a 260/280 ratio close to the ratio of the purified nanobody before biotinylation. In most cases this ratio should be between 0.5-0.7. Measure the final concentration and freeze aliquots in liquid nitrogen.

**CRITICAL STEP** Including later fractions with increased 260/280 ratios (>0.7) risks contaminating the final biotinylated ALFA nanobody preparations with excess free biotin that will compete with nanobody binding to streptavidin beads.

**PAUSE POINT** Frozen protein aliquots are stable at -80°C for years.

##### (Optional)#Assessing the biotinylation efficiency of the purified nanobodies ● Timing ∼4 h

The biotinylation efficiency of the purified nanobodies can be assessed by testing their binding to streptavidin beads. Use pre-chilled buffer and work on ice.

17 **Equilibrate beads:** Carefully resuspend magnetic Streptavidin beads until the slurry is homogenous and no clumps are left on the bottom or side of the bottle. Aliquot 25 µl slurry into a 1.5 ml tube and mix with 1 ml STB buffer. Retrieve beads by placing the tube into a magnetic rack, wait for 30 - 60 seconds to collect beads on the magnet and then aspirate all buffer.

18 **Prepare nanobodies:** Dilute 12 µg biotinylated nanobody in 60 µl STB buffer. Take an input sample by mixing 10 µl of the dilution with 2.5 µl 5x SDS-PAGE sample buffer.

**CRITICAL STEP** In order to assess biotinylation efficiency, it is essential to avoid oversaturating the beads. We found that the capacity of Pierce magnetic Streptavidin beads is around 0.5 µg of biotinylated nanobody fusion protein per 1 µl slurry of beads.

19 **Binding:** Remove tube from the magnetic rack and resuspend the beads with the remaining 50 µl diluted nanobody. Incubate for 30 min with occasional gentle flicking of the tube to prevent beads from settling.

20 Retrieve beads by placing the tube into a magnetic rack. Take off unbound fraction and take a sample by mixing 10 µl unbound fraction with 2.5 µl 5x SDS-PAGE sample buffer.

21 **Washing:** Wash beads by resuspending them in 1 ml STB buffer. Retrieve beads by placing the tube into a magnetic rack and then aspirate buffer. Repeat this step. For the last wash step, resuspend beads in 100 µl buffer to concentrate them in a smaller area. Retrieve beads and aspirate all buffer.

22 **Elution:** Resuspend beads in 60 µl 2x SDS-PAGE sample buffer containing 0.5 M urea and boil for 10 min. at 97°C. Retrieve beads and take off the elution to a new tube.

23 **SDS-PAGE analysis:** Analyze the test binding by SDS-PAGE and Coomassie staining. Load 12.5 µl of input and unbound samples, as well as 10 µl of the elution per lane. The expected outcome is shown in **Fig. 5b**.

? TROUBLESHOOTING

##### (Optional)#Assessing the activity of purified SENP^EuB^ ● Timing ∼3 h

24 Make up 10 µM (0.32 µg/µl) purified pTP396_His-Avi-SUMO^Eu^-anti-GFP nanobody in a final volume of 10 µl resuspension buffer with or without 250 nM purified SENP^EuB^.

25 Incubate both reactions for 20 min on ice

26 Add 10 µl 2x SDS-PAGE sample buffer to each sample. Analyze the test cleavage by SDS-PAGE and Coomassie staining. Load 10 µl per lane (∼1.6 µg of pTP396). The expected outcome is shown in **Fig. 5A**.

? TROUBLESHOOTING

##### Generation of a stable human suspension cell line ● Timing ∼8 d

#### Production of high-titer lentiviral supernatants ● Timing ∼5 d

Step 27 can be started in parallel to step 1.

27 **Take Lenti-X 293T cells into culture:** Thaw a single frozen Lenti-X 293T cell aliquot, containing around ∼8-10 x10^6^ cells and add to 10 ml DMEM in a 15 ml tube. Spin for 3 min. at 300 g to pellet cells and remove DMSO. Aspirate medium and resuspend cell pellet in 1 ml DMEM. Add drop-wise into a 15 cm dish containing 19 ml DMEM. Distribute cells evenly and incubate in a 37°C, 5% CO2 cell culture incubator.

**! CAUTION** Cell cultures are a potential biological hazard. Make sure to work in an approved laminar flow hood and use proper sterile technique. Adhere to the relevant institutional and governmental guidelines for recommended protective personal equipment and proper disposal of waste.

**CRITICAL STEP** Prevent Lenti-X 293T cells from reaching confluency. We typically split cells 1:10 every two days. Treat these cells very gently to preserve high transfection efficiency, which is essential to reach high lentiviral titer. Freeze aliquots of very early passages and avoid prolonged culture (> 1 month).

28 **Seed Lenti-X 293T cells into 6-well plates**: Once the 15 cm dish reaches ∼70% confluency, aspirate all culture medium, wash gently with 20 ml DPBS and detach cells by addition of 8 ml Trypsin-EDTA (0.25%). Incubate for 2 min at room temperature and then resuspend cells with 12 ml DMEM. Count resuspended cells and seed 1x10^6^ cells per 6-well in 2.5 ml DMEM. Incubate 6-well plate for ∼24 h.

29 (Optional) **Freeze aliquots**: Freeze excess resuspended HEK 293T cells from step 28. Pellet cells for 3 min at 300 g, aspirate medium and resuspend in DMEM containing 10 % (v/v) DMSO. Make 1 ml aliquots containing ∼8-10 x10^6^ cells. Place cell aliquots into a cell freezing container and store it at -80°C for 24 h. Transfer frozen cell aliquots into a cryo-tank for long-term storage.

30 **Transfection:** Proceed with transfection once cells reach ∼70-80% confluency to produce best lentiviral titers. Per 6-well mix 1.25 µg transfer plasmid, 0.94 µg psPAX2 and 0.32 µg pMD2.G in 250 µl Opti-MEM I Reduced-Serum Medium. Mix well and add 7.5 µl TransIT 293 Transfection Reagent. Mix again and incubate for 15 min at room temperature. Carefully add formed transfection complexes to 6-wells in drops and mix plate by swirling. **CRITICAL STEP** Transfer plasmids contain repetitive sequences and should be prepared from recombination-deficient (recA^-^) *E. coli* strains like Stellar.

**! CAUTION** Enhanced biosafety level measures are required for all following steps.

31 **Analyze transfection efficiency:** After 24-36 h analyze transfection efficiency of each 6- well by visually assessing the percentage of fluorescent cells on a tissue-culture microscope. Non-fluorescent wells usually indicate problems with transfection and will yield low lentiviral titers.

? TROUBLESHOOTING

32 **Harvest lentiviral supernatant:** 48 h after transfection harvest culture supernatant (∼2.5 ml) and spin for 5 min at 500 g in sterile 5 ml tubes. Use rotor buckets with aerosol-tight caps and assemble and disassemble them in the laminar flow hood. Use supernatant immediately to transduce Expi293F suspension cells (step 34).

**PAUSE POINT** Single-use aliquots of lentiviral supernatants can be snap-frozen in liquid nitrogen for storage at -80°C for multiple months with only slight loss of viral titer.

**CRITICAL** Repeated freeze-thaw cycles result in significant loss of lentiviral titer. It is best to not re-use once thawed aliquots.

#### Transduce and grow Expi293F suspension cells ● Timing ∼8 d

Step 33 can be started in parallel to step 1 and 27 to allow the Expi293F cells to recover for a few days before transduction.

33 **Take Expi293F cells into culture**: Thaw a 1 ml aliquot containing ∼20x10^6^ Expi293F cells and add to 10 ml Expi medium in a 15 ml tube. Spin for 3 min at 300 g to remove DMSO. Resuspend cell pellet in 1 ml Expi medium and transfer to 49 ml of the same medium in a 125 ml Erlenmeyer flask with vented cap.

**CRITICAL** Grow Expi293F cells in a 37°C orbital shaker with 8% CO2, >80% relative humidity and shaking at ∼125 rpm. Maintain Expi293F stock between 0.5-2x10^6^ cells/ml by diluting them 1:4 with fresh medium every two days. Freeze excess cells in Expi medium supplemented with 10% (v/v) DMSO to replenish cell aliquots.

**! CAUTION** Enhanced biosafety level measures are required for all following steps.

34 **Transduction with lentivirus**: In a new 125 ml Erlenmeyer flask with vented cap, mix 20 ml of Expi293F cells at 1x10^6^ cells/ml with 2.5 ml freshly harvested lentiviral supernatant. Then transfer flask to shaking incubator and grow overnight. If needed this step can easily be scaled up or down.

**CRITICAL STEP** Transduction efficiency depends strongly on lentiviral titer. This can be optimized by mixing a constant amount of suspension cells with increasing volume of lentiviral supernatant.

35 **Exchange medium:** The next day transfer culture into a 50 ml tube, place into rotor buckets with aerosol-tight caps in the laminar flow hood and then spin at 300 g for 3 min. Take off supernatant and mix with 10% bleach to inactivate lentivirus. Resuspend cell pellet in 50 ml Expi medium and transfer to a new 125 ml Erlenmeyer flask with vented cap. Grow cells for 72 h.

36 **Analyze transduction efficiency**: Analyze the ratio of fluorescent cells 48 h after transduction either visually using a tissue culture microscope or more quantitatively via flow cytometry. Protein expression can also be analyzed by Western blotting. If using inducible cells, take off 2 ml of culture to a 6-well plate and induce for 24 h with 1 µg/ml DOX before analysis of transduction efficiency as above.

? TROUBLESHOOTING

37 **Growth phase**: For small-scale expression trials, 50 ml is a good initial culture volume and cells can be easily be grown to around 8x10^6^cells/ml without any drop in viability if the expressed protein is not toxic.

**CRITICAL STEP** If using inducible cells make sure to add 1 µg/ml DOX to the culture medium. Induction time can be varied, but 24-48 h before harvest is a good starting point. Prior to induction, make up a separate stock of non-induced cells at 0.5x10^6^ cells/ml in 50 mL in a new flask to keep them in culture.

38 **Cell harvest:** Remove flask from shaking incubator. If using non-inducible cells, count cells and make up a separate stock at around 0.5x10^6^cells/ml in 50 ml in a new flask to keep them in culture in case sorting is necessary or the culture needs to be expanded for large-scale preparations. Transfer remaining cells into a 50 ml tube and pellet cells as described above. Wash cells by resuspending them in 50 ml DPBS and repeating the spin. Pour off liquid, weigh dry cell pellet and proceed directly to cell lysis.

**PAUSE POINT** Alternatively, cell pellets can be frozen in liquid nitrogen and stored at -80°C for multiple months.

##### (Optional)#Sorting of transduced Expi293F cells ● Timing ∼7-9 d

39 **Harvest cells:** Harvest 45x10^6^ transduced and non-transduced Expi293F cells as described above. Resuspend the washed cell pellets in 3 ml DPBS supplemented with 20% (v/v) FBS to a concentration of ∼15x10^6^ cells/ml. Filter resuspended cells through strainer caps into 5 ml round bottom FACS tubes.

40 **Adjust gates:** Analyze non-transduced Expi293F cells on a SONY SH800S or equivalent cell sorter and adjust gates to select alive and single cells, as well as to determine background fluorescence level. Set gate for transduced fluorescent cells accordingly.

41 **Sort cells:** Aim to collect at least 2-4x10^6^ cells in a 15 ml collection tube pre-filled with 3 ml Expi medium containing 0.5x Pen-Strep. Pellet collected cells to remove sheath fluid and resuspend in 10 ml Expi medium containing 0.5x Pen-Strep. Transfer to a new 125 ml Erlenmeyer flask with vented cap and grow overnight.

**CRITICAL STEP** Since most cell sorters are not operated under perfectly sterile conditions, it is essential to add Pen-Strep to prevent contamination. However, only 0.5x Pen-Strep is used since Expi293F cells are more sensitive to antibiotics than regular HEK 293T cells.

42 **Exchange medium:** 24 h after sorting, pellet cells and aspirate medium. Resuspend in 10 ml Expi medium without Pen-Strep and transfer back to the same flask.

#### **CRITICAL STEP** Prolonged exposure to Pen-Strep reduces cell viability

43 **Recovery**: Let cells recover by growing them for 5-7 days. Assess cell density every day, replenish medium and dilute cells to grow up a new stock of sorted cells. Freeze aliquots of early passages.

44 **Cell lysis** ● Timing **∼2 h**

**A) Mechanical lysis without detergent**

**CRITICAL STEP** Lysis buffer composition is protein-specific and should be optimized in small-scale experiments.

(1) **Resuspension**: Resuspend cell pellet in pre-chilled lysis buffer. Frozen cell pellets should be thawed quickly in a lukewarm water bath in the presence of lysis buffer and removed to ice immediately once thawing nears completion.
(2) **Lysis**: Lyse cells with multiple passes in dounce homogenizer using a tight fit pestle. Alternatively, you may also use a motor-driven potter-elvehjem homogenizer. Monitor the progression of cell lysis on a cell counter or tissue-culture microscope using trypan blue exclusion. Continue lysis until no more intact cells remain.
(3) **Remove cell debris:** Centrifuge cell lysate for 30 min at 35,000 g and 4°C. Take off supernatant and proceed to affinity purification.

**B) Lysis in detergent**

**CRITICAL STEP** Optimal detergent type, salt concentration, as well as ratio of solubilization buffer to cell pellet weight is membrane protein, as well as application-specific and should be optimized in small-scale experiments. Gentle detergents like GDN are strongly preferred to maintain protein complex stability and to preserve transient protein interactions over harsher detergents like Triton-X-100 or NP-40, which if tolerated enable purifications with higher purity. We routinely solubilize whole cell pellets, but in certain cases it is advantageous to prepare a membrane fraction prior to solubilization. The steps below offer a good starting point.

(1) **Solubilization:** Add 7 ml pre-chilled solubilization buffer, containing 1% (w/v) of a detergent of choice, per each gram of cell pellet and resuspend. Frozen cell pellets should be thawed quickly in a lukewarm water bath in the presence of solubilization buffer and removed to ice immediately once thawing nears completion. Incubate mixing head-over-tail for 30 min at 4°C.
(2) **Remove cell debris:** Centrifuge cell lysate for 30 min at 35,000 g and 4°C. Take off supernatant and proceed to affinity purification.

##### Affinity purification ● Timing ∼3 h

**CRITICAL STEP** The amount of beads and nanobody needed depend on the expression level of the protein of interest and should be optimized in small-scale experiments. The steps below offer a good starting point. Use pre-chilled buffer and work on ice.

45 **Equilibrate beads:** For every gram of cell pellet, equilibrate 60 µl of magnetic streptavidin beads slurry. Carefully resuspend beads until the slurry is homogenous and no clumps are left on the bottom or side of the bottle. Aliquot slurry into a 1.5 ml tube. Retrieve beads by placing the tube into a magnetic rack, wait for 30 sec to 1 min to collect all beads and then aspirate all liquid. Resuspend in 1 ml wash buffer, retrieve beads as above and aspirate wash buffer.

**CRITICAL STEP** For membrane proteins a detergent at a concentration above its critical micelle concentration (CMC) needs to be included in wash and elution buffers to keep it solubilized. We recommend using a concentration that corresponds to 2.5x CMC.

46 **Immobilize nanobody:** For every 60 µl of magnetic Streptavidin beads slurry, immobilize 20 µg of biotinylated GFP or ALFA nanobody. Resuspend beads in 500 µl wash buffer containing pre-diluted biotinylated nanobody and incubate beads for 15 min rotating head-over-tail at 4°C.

47 **Block beads**: Retrieve beads using a magnet and aspirate wash buffer. Remove from the magnet and resuspend in 500 µl wash buffer containing 100 µM biotin or dPEG24-biotin acid. Incubate for 5 min on ice to block unoccupied biotin binding sites.

**CRITICAL STEP** Blocking with biotin strongly reduces background binding of endogenous biotinylated proteins. Blocking with dPEG24-biotin acid adds additional negative charge and further reduces non-specific background binding.

48 **Incubate with lysate:** Retrieve beads using magnetic rack and aspirate blocking buffer. Resuspend beads with cleared cell lysate and mix rapidly. Incubate the mixture for 1 h at 4°C rotating.

49 **Washing:** Retrieve beads on magnet. Depending on the final scale a magnetic rack that can hold 15 ml or even 50 ml tubes should be used at this step. Aspirate cell lysate and transfer all beads into a 1.5 ml tube using 1 ml wash buffer. Wash beads three times with 1 ml wash buffer. For the fourth wash, resuspend beads with 100 µl wash buffer (-ATP) and transfer to a new tube. Use slightly more volume if significantly more beads were used.

50 **Elution:** Resuspend beads in wash buffer containing 250 nM purified SENP^EuB^. Use a volume corresponding to the original volume of bead slurry used. To keep the eluate more concentrated, up to 1/3^rd^ of the original volume can be used. Incubate for 20 min on ice.

**CRITICAL STEP** If a soluble protein is stable in the presence of detergents (like 0.05% [v/v] Tween-20 or Triton-X-100), these can be included during elution to improve recovery by preventing non-specific binding of the cleaved protein to the beads.

51 (Optional) Spin eluate for 5 min at 15,000 g at 4°C to pellet magnetic beads that sometimes get carried over. Take off supernatant.

52 **Post-Elution**: Resuspend beads in 2x SDS-PAGE sample buffer containing 0.5 M urea and boil for 10 min at 97°C. Retrieve beads on magnet and take off post-elution sample.

53 **SDS-PAGE analysis:** Analyze samples of the eluate and post-elution by SDS-PAGE. We frequently load 1, 2 and 4 µl per lane and stain the gel with either Sypro Ruby or Coomassie.

? TROUBLESHOOTING

##### (Optional)#Size-exclusion chromatography ● Timing 1 d

An additional size-exclusion chromatography run might be useful to remove excess tagged subunit and nanobody. We prefer the Superose 6 Increase 3.2/300 column due its small column volume of only 2.4 ml, which limits sample dilution during the run.

54 **Column equilibration**: Equilibrate HPLC, Superose 6 Increase 3.2/300 column and 50 µl sample loop in filtered and degassed wash buffer without protease inhibitor cocktail.

55 **Sample loading**: Load sample into a 50 µL hamilton syringe and inject into sample loop. **CRITICAL STEP** Spin eluate for 10 min at 15,000 g at 4°C before loading to pellet magnetic beads and aggregated protein. Take off supernatant. If eluate volume is above 50 µl concentrate using ultrafiltration spin columns with the appropriate molecular weight cut-off.

56 **Elution**: Inject sample loop content onto column and elute in 100 µl fractions over 1.2x column volume. Measure absorbance at 280 nm, optionally at 490 nm to detect GFP fluorescence or 260 nm to detect non-protein contaminants.

57 **SDS-PAGE analysis**: Choose fractions spanning sample peak and prepare SDS-PAGE sample by mixing 10 µl of each fraction with 2.5 µl 5x SDS-PAGE sample buffer. If this is a high yield purification you may instead prefer to dilute a smaller volume of each fraction to 10 µL with wash buffer to avoid overloading the gel. Analyze samples by SDS-PAGE and stain with either Sypro Ruby or Coomassie.

58 Wash HPLC, Superose 6 Increase 3.2/300 column and 50 µl sample loop with degassed ddH2O.

59 Pool protein-containing fractions. If necessary, concentrate protein using ultrafiltration spin columns with the appropriate molecular weight cut-off.

### Timing

Generation of biotinylated nanobodies and SENP^EuB^: ∼4-5 d Steps 1-12, Preparation of biotinylated GFP nanobodies: ∼4 d

Steps 1-12, Preparation of SENP^EuB^ protease and biotin ligase BirA: ∼4 d

Steps 1-12, Preparation of ALFA nanobodies: ∼4 d

Steps 13-16, *In vitro* biotinylation of ALFA nanobodies with purified BirA: ∼5 h

Steps 17-23, (Optional) Assessing the biotinylation efficiency of the purified nanobodies: ∼4 h

Steps 24-26, (Optional) Assessing the activity of purified SENP^EuB^: ∼3 h

#### Generation of a stable human suspension cell line: ∼8 d

Steps 27-32, Production of high-titer lentiviral supernatants: ∼5 d Steps 33-38, Transduce and grow Expi293F suspension cells: ∼8 d Steps 39-43, (Optional) Sorting of transduced Expi293F cells: ∼7-9 d Step 44, Cell lysis: ∼2 h

Steps 45-53, Affinity purification: ∼3 h

Steps 54-59, (Optional) Size-exclusion chromatography: ∼1 d

### Troubleshooting

Troubleshooting advice is listed in Table 3.

**Table 3.** Troubleshooting table

| Step | Problem | Possible reason | Solution |
| --- | --- | --- | --- |
| 11 | No protein after Ni<sup>2+</sup>-purification | Forgot to add IPTG to induce protein expression | Add IPTG to 0.2 mM |
| 11 | No protein after Ni<sup>2+</sup>-purification | Forgot to add imidazole to elution buffer | Use elution buffer containing imidazole at 500 mM |
| 11 | Low protein yield after Ni<sup>2+</sup>-purification | Protein might have aggregated during expression due to elevated expression temperature | Make sure to adapt diluted main culture to 18°C for 1 h before induction and maintain induced main culture at 18°C. |
| 11 | Low protein yield after Ni<sup>2+</sup>-purification | Protein might have aggregated during purification | Keep *E. coli* cells/ lysate/ purified protein on ice at all times and prevent foaming during sonication |
| 11 | *E. coli* purified protein is partially degraded | Activation of *E. coli* proteases during cell lysis | Make sure to add 1 mM PMSF to resuspension buffer and keep *E. coli* cells/ lysate/ purified protein on ice at all times |
| 23 | GFP Nanobodies produced in *E. coli* CVB101 do not bind to Streptavidin beads | Forgot to supplement CVB101 main culture with biotin or forgot to use chloramphenicol to maintain BirA plasmid | Make sure to propagate CVB101 with 10 µg/ml chloramphenicol and add 50 µM biotin to main culture |
| 23 | *In vitro* biotinylated ALFA nanobodies do not bind to Streptavidin beads | Poor quality BirA preparation or wrong biotinylation buffer composition | Check BirA by SDS-PAGE and check 5x biotinylation buffer composition. Repeat BirA preparation. |
| 26 | SENP<sup>EuB</sup> does not cleave nanobodies in solution | Poor quality SENP<sup>EuB</sup> preparation | Check SENP<sup>EuB</sup> by SDS-PAGE. Repeat SENP<sup>EuB</sup> preparation. |
| 31 | Transfected Lenti-X 293T cells non-fluorescent | Poor DNA quality | Check plasmid DNA integrity and prepare new stock |
| 36 | Low transduction efficiency | Lenti-X 293T cells passaged for prolonged time and/or overgrown prior to seeding (bad cell health) | Thaw a new Lenti-X 293T cell aliquot and avoid overgrowing and passaging them > 1 month |
| 36 | Low transduction efficiency | Transfected Lenti-X 293T cells at too high or too low confluency | Make sure to transfect at around 70-80% confluency |
| 36 | Low transduction efficiency | Forgot or added wrong amounts or bad quality of packaging and envelope plasmids | Make sure to add correct amounts of packaging and envelope plasmids and consider preparing new stocks |
| 36 | Low transduction efficiency | DNA insert size comes close to or exceeds packaging limit (~8 kbp between 5'-3' LTRs) | Sort transduced cells and grow up a fully transduced population |
| 36 | Low transduction efficiency | Protein of interest is toxic to cells | Try expressing an inactive mutant, or use an alternative expression method such as chemical transfection or recombinase-mediated cell line generation |
| 53 | Low protein yield from Expi293F cells | Poor lentiviral titer or transduction efficiency | Repeat lentiviral preparation or sort transduced cells and grow up a fully transduced population |
| 53 | Low protein yield from Expi293F cells | Inefficient protease elution | Analyze post-elution sample by SDS-PAGE and check for uncleaved bait (TagON) or bait-prey complex (TagOFF). Check protease activity in solution (see steps 24-26) and repeat protease preparation if inactive. In some cases the SUMO tag may be less accessible, requiring increased protease concentration and incubation time to achieve efficient elution. |
| 53 | Low protein yield from Expi293F cells | Eluted protein non-specifically sticks to beads (sometimes for soluble proteins in buffer without detergent) | Include low concentration of detergent (e.g. 0.05 % [v/v] Tween-20 or Triton-X-100) in elution buffer to reduce sticking. Block Streptavidin beads with biotin-PEG-COOH instead of biotin to increase charge repulsion between beads and protein. The cleaved protein may be released in additional wash steps. |
| 53 | Low protein yield from Expi293F cells | Protein aggregated due to suboptimal lysis buffer composition | Optimize lysis buffer composition e.g. salt or detergent concentration in small-scale pre-tests. |
| 53 | Low recovery of the prey's endogenous protein complex subunits | Tagged protein is stable on its own after overexpression | Consider tagging a different, less stable subunit of the complex. Alternatively: 1) Use Freestyle 293-F cells instead. 2) Use inducible Expi293F cells line and vary DOX induction time. 3) Exchange CMV to weaker PGK or UbC promoter or remove WPRE element. 4) Purify membrane proteins from the membrane fraction rather than whole cells. |

### Anticipated results

We have used this protocol as outlined in **Fig. 1** to isolate numerous soluble and membrane-bound proteins for structural, functional, and mass spectrometry analysis (**Fig. 3**). For proteins that are not part of a protein complex, high quality preparations can easily be achieved with often only minimal prior optimization. In certain cases, lysis buffer composition needs to be optimized in small-scale purification trials to obtain best results. In cases where little prior knowledge exists about a protein complex of interest, it is first necessary to optimize which subunit of the complex to tag and also at which terminus to tag this subunit.

Another critical consideration for the success of the protocol is to take great care to achieve high-titer lentiviral production as outlined above. Only with good lentiviral preparations can high transduction efficiencies of suspension cells be reached. In many cases, transduction efficiencies are readily between 80-90%, e.g. for EMC3-GFP (**Fig. 6**). In certain cases, e.g. for much larger proteins, lower transduction efficiencies necessitate either larger-scale cultures to achieve a comparable yield or an additional sorting step to obtain a fully transduced polyclonal cell line. Sorting typically postpones first purification trials by one extra week, which is required for recovery and expansion of the sorted cells.

**Fig. 6.**
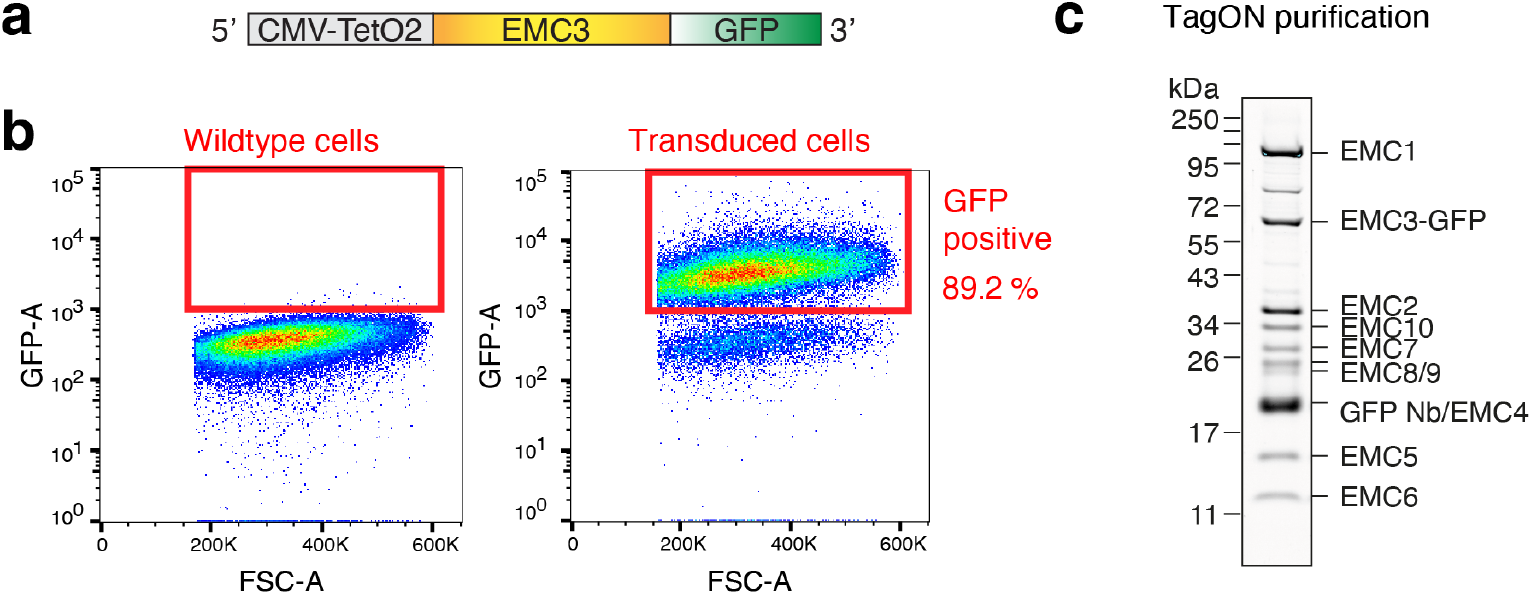
TagON purification of the EMC from a stable Expi293F EMC3-GFP suspension cell line. (**a**) Schematic overview of the EMC3-GFP expression cassette encoded on the lentiviral transfer plasmid. (**b**) Flow cytometry analysis of Expi293F cells 48 h after transduction as described in steps 33-38. Cells were gated first for alive cells (SSC-A vs. FSC-A), second for single cells (FSC-A vs. FSC-H) and third for GFP-fluorescent cells (GFP-A vs. FSC-A). Nearly 90% of cells were GFP positive. (**c**) TagON purification of the EMC via EMC3-GFP. An aliquot of the elution was analyzed by SDS-PAGE and Sypro Ruby staining.

Using this protocol, we have achieved improved yields for challenging multi-subunit membrane protein complexes. For example, we could isolate 0.45 mg of EMC or ∼0.8 mg of MTCH2 per 1 L of suspension cell culture. Much greater yields can be achieved for soluble protein complexes and in particular for soluble monomeric proteins.

## Acknowledgments

We thank Pamela Bjorkman for access to her lab’s cell sorter, as well as the Caltech Flow Cytometry facility. This work was supported by: the Heritage Medical Research Institute (RMV), the NIH’s National Institute Of General Medical Sciences DP2GM137412 (RMV), the Deutsche Forschungsgemeinschaft (TP), and the Tianqiao and Chrissy Chen Institute (TP, MH).

## Data availability statement

The lentiviral transfer plasmids and bacterial expression plasmids described in this study are available from Addgene. Addgene IDs of all plasmids are listed in Table 1.

## List of supplementary information

Supplementary Data 1. A cloning guide for the pTS93-116 lentiviral transfer plasmid toolbox.

Supplementary Data 2. E. coli protein expression data sheet.

## Supplementary Information

**Extended Data Figure 1.**
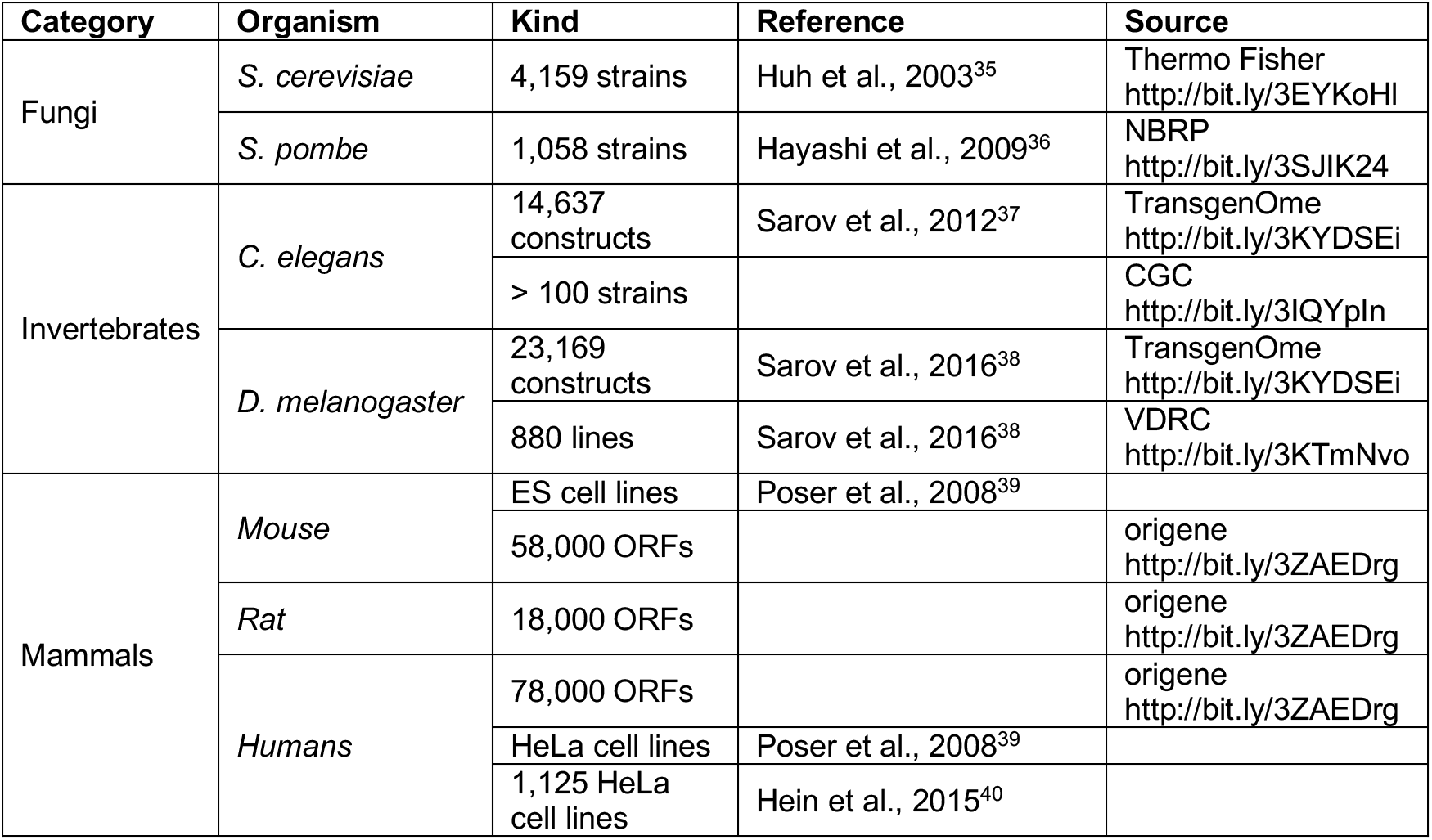
Publicly or commercially available GFP-tagged plasmids, cell lines or transgenic organisms. Thousands of plasmids encoding GFP- and ALFA-tagged proteins from various organisms can also be obtained from Addgene. ORF = open-reading frame. ES = embryonic stem cell.

**Extended Data Figure 2.**
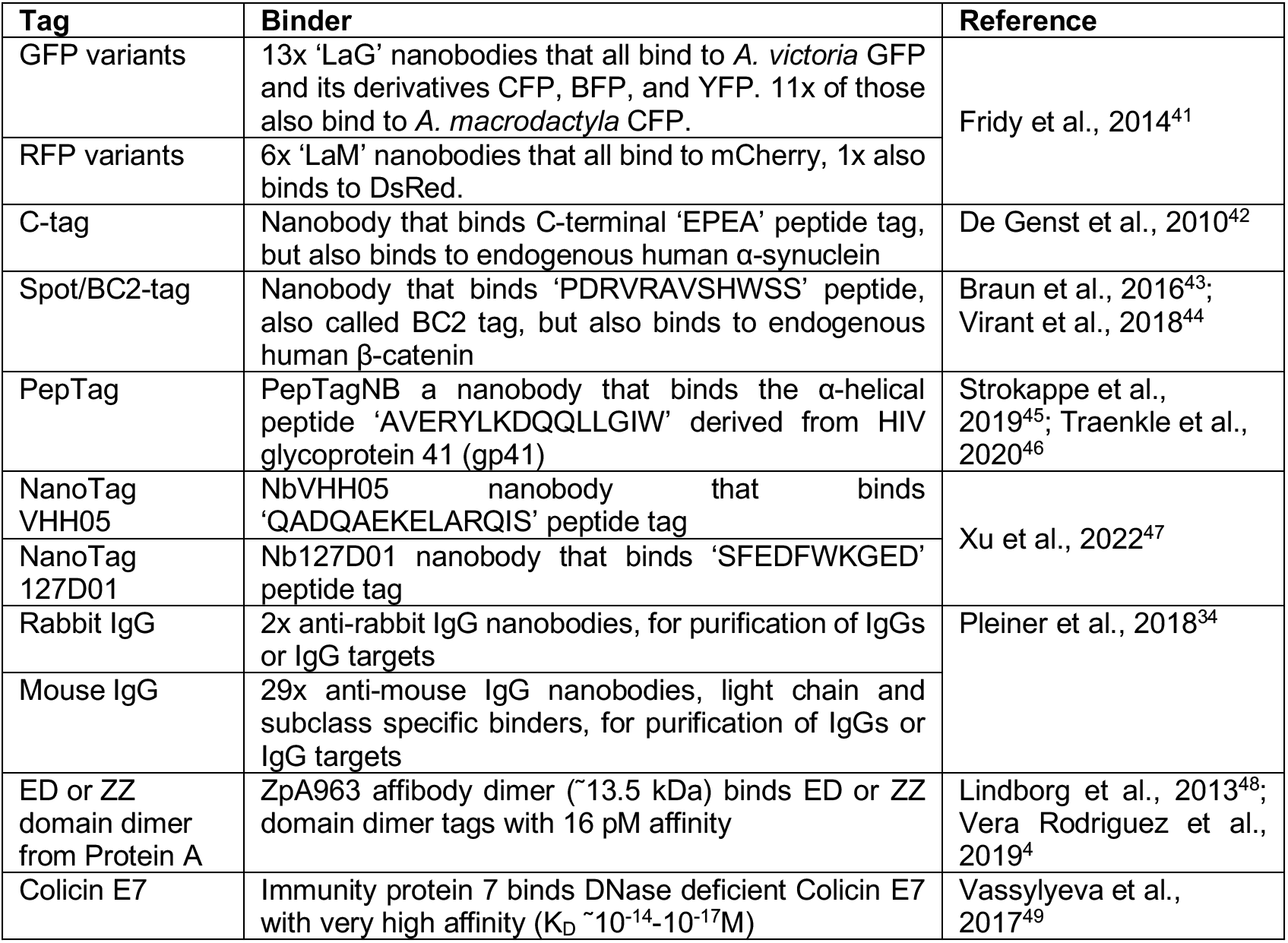
Selection of previously characterized affinity binder pairs that could be used for TagON/OFF purifications.

**Extended Data Figure 3.**
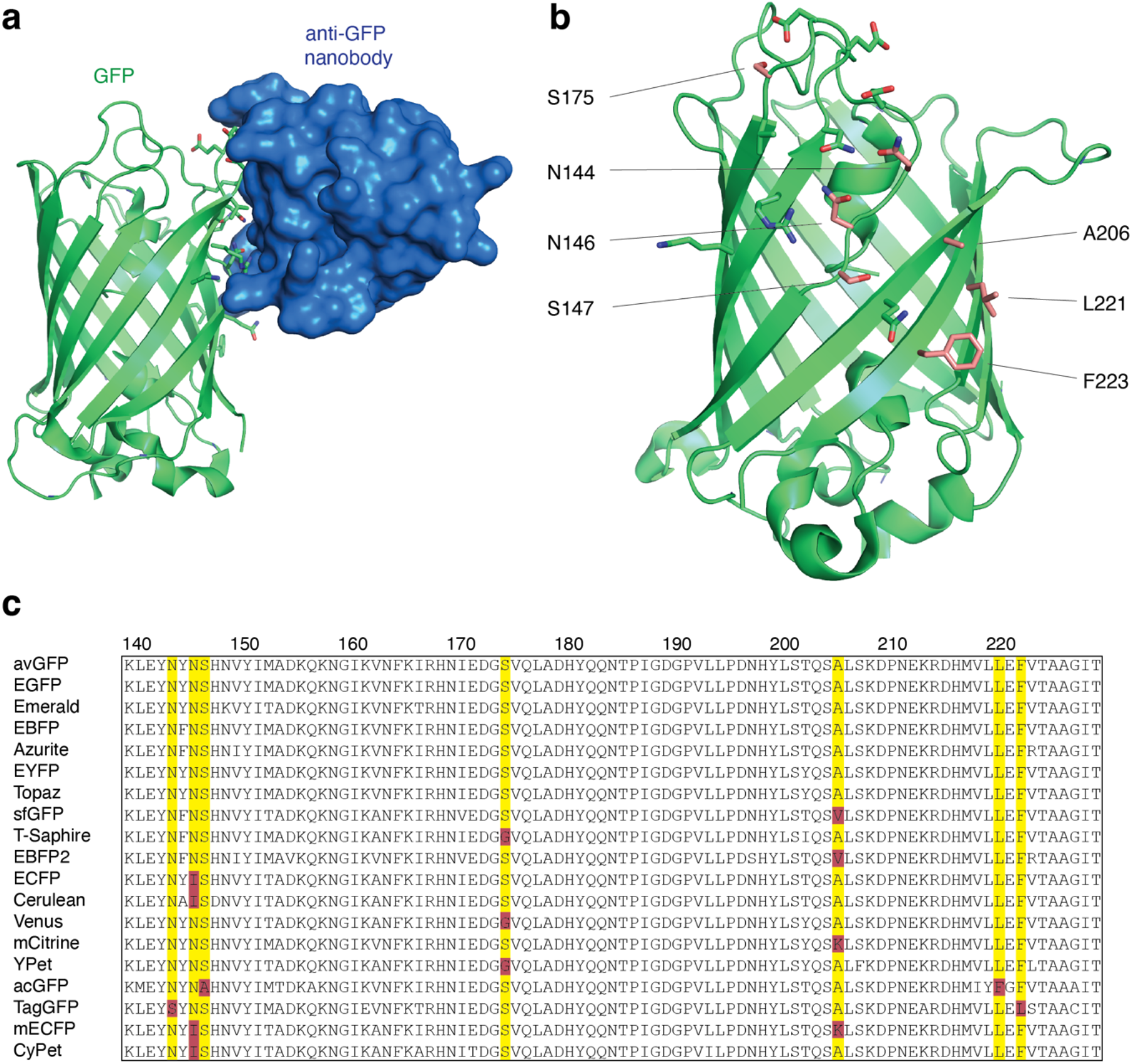
Overview of anti-GFP nanobody compatible fluorescent protein variants. (**a**) Crystal structure of GFP bound to anti-GFP nanobody (Nb) (PDB ID: 3K1K)^25^ with GFP shown in green with cartoon rendering, anti-GFP Nb shown in blue with surface rendering, and specific residues on GFP that make contact with the anti-GFP Nb shown in stick rendering. (**b**) Front-view of the anti-GFP nanobody binding surface of GFP with participating residues shown in stick rendering. Residues mutated in other fluorescent protein variants colored in salmon. (**c**) Multiple sequence alignment of various fluorescent protein variants. Columns corresponding to residues contacted by the anti-GFP Nb are highlighted in yellow, and any mutations to these positions are highlighted in red. Mutation of I146N was previously shown to restore anti-GFP Nb binding in CFP variants^50^.

**Extended Data Figure 4.**
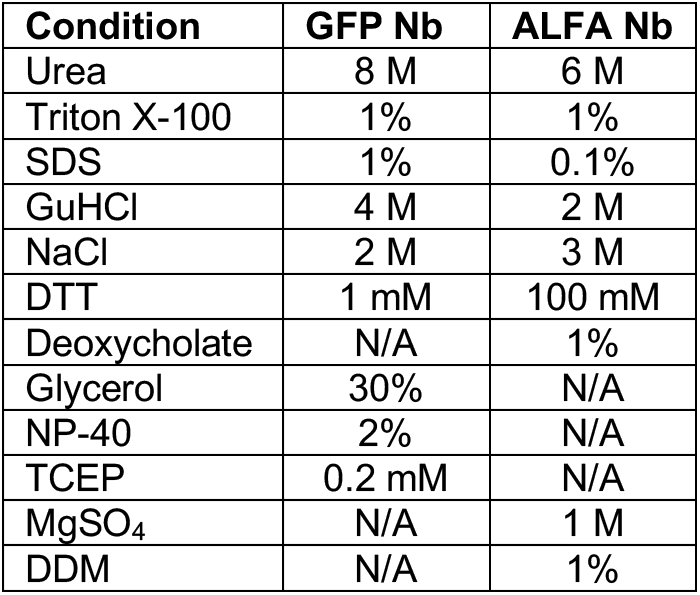
Anti-GFP and anti-ALFA nanobodies withstand harsh buffer conditions. Comparison of buffer conditions tolerated by the anti-GFP nanobody Enhancer (https://bit.ly/3kLbpHr) and the anti-ALFA^ST^ nanobody^3^. Nb = nanobody; N/A = data not available.

## References

1. Elegheert, J. et al. Lentiviral transduction of mammalian cells for fast, scalable and high-level production of soluble and membrane proteins. Nat Protoc 13, 2991–3017 (2018).

2. Pleiner, T. et al. Structural basis for membrane insertion by the human ER membrane protein complex. Science 369, 433–436 (2020).

3. Götzke, H. et al. The ALFA-tag is a highly versatile tool for nanobody-based bioscience applications. Nat Commun 10, 4403 (2019).

4. Vera Rodriguez, A., Frey, S. & Görlich, D. Engineered SUMO/protease system identifies Pdr6 as a bidirectional nuclear transport receptor. J Cell Biol 218, 2006–2020 (2019).

5. Pleiner, T. et al. Nanobodies: site-specific labeling for super-resolution imaging, rapid epitope-mapping and native protein complex isolation. Elife 4, e11349 (2015).

6. Aksu, M. et al. Xpo7 is a broad-spectrum exportin and a nuclear import receptor. J Cell Biol 217, 2329–2340 (2018).

7. Chen, J. et al. Structure of an endogenous mycobacterial MCE lipid transporter. bioRxiv doi: 10.1101/2022.12.08.519548 (2022).

8. Juszkiewicz, S. & Hegde, R. S. Quality Control of Orphaned Proteins. Mol Cell 71, 443–457 (2018).

9. Pleiner, T. et al. WNK1 is an assembly factor for the human ER membrane protein complex. Mol Cell 81, 2693–2704.e12 (2021).

10. Guna, A. et al. MTCH2 is a mitochondrial outer membrane protein insertase. Science 378, 317–322 (2022).

11. Kawate, T. & Gouaux, E. Fluorescence-detection size-exclusion chromatography for precrystallization screening of integral membrane proteins. Structure 14, 673–681 (2006).

12. Voorhees, R. M. & Hegde, R. S. Structures of the scanning and engaged states of the mammalian SRP-ribosome complex. Elife 4, e07975 (2015).

13. McShane, E. et al. Kinetic Analysis of Protein Stability Reveals Age-Dependent Degradation. Cell 167, 803–815.e21 (2016).

14. de Felipe, P. et al. E unum pluribus: multiple proteins from a self-processing polyprotein. Trends Biotechnol 24, 68–75 (2006).

15. Goehring, A. et al. Screening and large-scale expression of membrane proteins in mammalian cells for structural studies. Nat Protoc 9, 2574–2585 (2014).

16. Chaudhary, S., Pak, J. E., Gruswitz, F., Sharma, V. & Stroud, R. M. Overexpressing human membrane proteins in stably transfected and clonal human embryonic kidney 293S cells. Nat Protoc 7, 453–466 (2012).

17. O’Gorman, S., Fox, D. T. & Wahl, G. M. Recombinase-mediated gene activation and site-specific integration in mammalian cells. Science 251, 1351–1355 (1991).

18. Leonetti, M. D., Sekine, S., Kamiyama, D., Weissman, J. S. & Huang, B. A scalable strategy for high-throughput GFP tagging of endogenous human proteins. Proc Natl Acad Sci U S A 113, E3501–E3508 (2016).

19. Koch, B. et al. Generation and validation of homozygous fluorescent knock-in cells using CRISPR– Cas9 genome editing. Nat Protoc 13, 1465–1487 (2018).

20. Cho, N. H. et al. OpenCell: Endogenous tagging for the cartography of human cellular organization. Science 375, eabi6983 (2022).

21. Nilsen, T. W. Preparation of nuclear extracts from HeLa cells. Cold Spring Harbor Protocols 2013, pdb. prot075176 (2013).

22. Seddon, A. M., Curnow, P. & Booth, P. J. Membrane proteins, lipids and detergents: not just a soap opera. Biochimica et Biophysica Acta (BBA)-Biomembranes 1666, 105–117 (2004).

23. Lee, S. C. et al. A method for detergent-free isolation of membrane proteins in their local lipid environment. Nat Protoc 11, 1149–1162 (2016).

24. Rothbauer, U. et al. Targeting and tracing antigens in live cells with fluorescent nanobodies. Nat Methods 3, 887–889 (2006).

25. Kirchhofer, A. et al. Modulation of protein properties in living cells using nanobodies. Nat Struct Mol Biol 17, 133–138 (2010).

26. Pédelacq, J.-D., Cabantous, S., Tran, T., Terwilliger, T. C. & Waldo, G. S. Engineering and characterization of a superfolder green fluorescent protein. Nature biotechnology 24, 79–88 (2006).

27. Schatz, P. J. Use of peptide libraries to map the substrate specificity of a peptide-modifying enzyme: a 13 residue consensus peptide specifies biotinylation in Escherichia coli. Biotechnology (N Y) 11, 1138–1143 (1993).

28. Beckett, D., Kovaleva, E. & Schatz, P. J. A minimal peptide substrate in biotin holoenzyme synthetase-catalyzed biotinylation. Protein Sci 8, 921–929 (1999).

29. Fairhead, M. & Howarth, M. Site-specific biotinylation of purified proteins using BirA. Methods Mol Biol 1266, 171–184 (2015).

30. Frey, S. & Görlich, D. The Xenopus laevis Atg4B protease: Insights into substrate recognition and application for tag removal from proteins expressed in pro-and eukaryotic hosts. PLoS One 10, e0125099 (2015).

31. Liu, L., Spurrier, J., Butt, T. R. & Strickler, J. E. Enhanced protein expression in the baculovirus/insect cell system using engineered SUMO fusions. Protein Expr Purif 62, 21–28 (2008).

32. Qin, J. Y. et al. Systematic comparison of constitutive promoters and the doxycycline-inducible promoter. PLoS One 5, e10611 (2010).

33. Frey, S. & Görlich, D. A new set of highly efficient, tag-cleaving proteases for purifying recombinant proteins. J Chromatogr A 1337, 95–105 (2014).

34. Pleiner, T., Bates, M. & Görlich, D. A toolbox of anti-mouse and anti-rabbit IgG secondary nanobodies. J Cell Biol 217, 1143–1154 (2018).

35. Huh, W.-K. et al. Global analysis of protein localization in budding yeast. Nature 425, 686–691 (2003).

36. Hayashi, A. et al. Localization of gene products using a chromosomally tagged GFP-fusion library in the fission yeast Schizosaccharomyces pombe. Genes to cells 14, 217–225 (2009).

37. Sarov, M. et al. A genome-scale resource for in vivo tag-based protein function exploration in C. elegans. Cell 150, 855–866 (2012).

38. Sarov, M. et al. A genome-wide resource for the analysis of protein localisation in Drosophila. Elife 5, e12068 (2016).

39. Poser, I. et al. BAC TransgeneOmics: a high-throughput method for exploration of protein function in mammals. Nat Methods 5, 409–415 (2008).

40. Hein, M. Y. et al. A human interactome in three quantitative dimensions organized by stoichiometries and abundances. Cell 163, 712–723 (2015).

41. Fridy, P. C. et al. A robust pipeline for rapid production of versatile nanobody repertoires. Nat Methods 11, 1253–1260 (2014).

42. De Genst, E. J. et al. Structure and properties of a complex of α-synuclein and a single-domain camelid antibody. Journal of Molecular Biology 402, 326–343 (2010).

43. Braun, M. B. et al. Peptides in headlock–a novel high-affinity and versatile peptide-binding nanobody for proteomics and microscopy. Scientific reports 6, 19211 (2016).

44. Virant, D. et al. A peptide tag-specific nanobody enables high-quality labeling for dSTORM imaging. Nature communications 9, 930 (2018).

45. Strokappe, N. M. et al. Super potent bispecific llama VHH antibodies neutralize HIV via a combination of gp41 and gp120 epitopes. Antibodies 8, 38 (2019).

46. Traenkle, B., Segan, S., Fagbadebo, F. O., Kaiser, P. D. & Rothbauer, U. A novel epitope tagging system to visualize and monitor antigens in live cells with chromobodies. Scientific reports 10, 14267 (2020).

47. Xu, J. et al. Protein visualization and manipulation in Drosophila through the use of epitope tags recognized by nanobodies. Elife 11, e74326 (2022).

48. Lindborg, M. et al. High-affinity binding to staphylococcal protein A by an engineered dimeric Affibody molecule. Protein Engineering, Design & Selection 26, 635–644 (2013).

49. Vassylyeva, M. N. et al. Efficient, ultra-high-affinity chromatography in a one-step purification of complex proteins. Proc Natl Acad Sci U S A 114, E5138–E5147 (2017).

50. Kubala, M. H., Kovtun, O., Alexandrov, K. & Collins, B. M. Structural and thermodynamic analysis of the GFP: GFP-nanobody complex. Protein Sci 19, 2389–2401 (2010).

