## Supplementary Data 1 for "A nanobody-based strategy for rapid and scalable purification of native human protein complexes"

### Cloning guide for the pTS93-116 lentiviral transfer plasmid toolbox

The pTS93-116 lentiviral transfer plasmid toolbox was created to allow easy implementation of our anti-GFP/ALFA nanobody-based purification strategy. It contains plasmids that allow fusion of a protein of interest (POI) to either N- or C-terminal GFP or ALFA peptide tags with or without additional protease cleavage sites. This guide describes how to clone your POI into these plasmids to create a transfer plasmid that can be co-transfected with the packaging plasmids psPAX2 (Addgene #12260) and pMD2.G (Addgene #12259) into Lenti-X 293T cells to create lentivirus. The resulting lentiviral supernatant can then be used to transduce for example human Expi 293F suspension cells to express your tagged POI for purification trials.

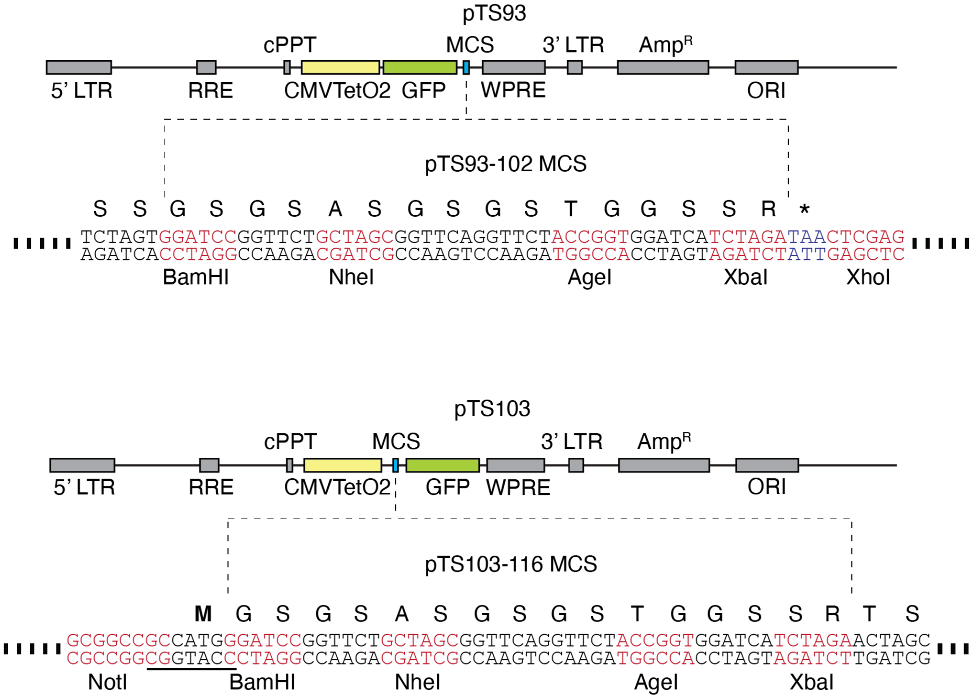


**Figure 1.** (Top) Schematic of pTS93, with detailed sequence view of the multiple cloning site (MCS) for pTS93-pTS102. (Bottom) Schematic of pTS103, with detailed sequence view of the MCS for pTS103-116 (Kozak sequence underlined). 5' LTR = 5' long terminal repeat, RRE = Rev response element, cPPT = central polypurine tract, CMVTetO2 = cytomegalovirus promoter with 2x TetO elements, MCS = multiple cloning site, WPRE = Woodchuck hepatitis virus post-transcriptional regulatory element, 3' LTR = 3' long terminal repeat, Amp$^R$ = Beta-lactamase (ampicillin/carbenicillin resistance), ORI = origin of replication

All transfer plasmids contain the same multiple cloning site (MCS) with 4 unique restriction sites (**Fig. 1**). Using the same restriction site, multiple plasmids encoding the POI fused to either GFP or ALFA tag at N- or C-terminus can therefore easily be generated during initial optimization trials. Besides restriction enzyme digest, we routinely clone POIs into these plasmids using Gibson assembly. To choose a method, consider the following:

| | Restriction enzyme cloning | Gibson cloning |
|---|---|---|
| Template DNA features | Insert sequence needs to lack the chosen MCS restriction sites | Internal restriction sites in insert don't interfere |

| | | |
|---|---|---|
| Modularity | Modular, digested insert fragment can be ligated into any plasmid | Not modular, different primers needed to clone insert into different backbones |
| Speed | Restriction digest adds extra step after PCR | There's no need to digest the PCR product |
| Primer length | Requires shorter primer overhangs (6 bp flanking + 6 bp restriction site) | Requires longer primer overhangs (15-40 bp Gibson overhangs) |

General plasmid design considerations:

- Make sure to include a stop codon in your reverse primer when cloning into pTS93-pTS102. There is a stop codon after the MCS but at least 2 amino acids will be added if no stop codon is added to the insert fragment.
- When adding an N-terminal tag (using pTS93-pTS102) check the Uniprot annotation for your POI to see if the initiator methionine is removed. If so, it is best to avoid including this codon in your insert.
- When adding a C-terminal tag, the N-terminus of your POI will include additional amino acids from the MCS sequence as well as an initiator methionine that we have included in the backbone to ensure a functional Kozak consensus sequence (5'-(gcc)gccRccATGG-3').
- If it is important to maintain the endogenous N-terminus, you can digest your backbone with NotI, but you must ensure that the resulting sequence contains a functional Kozak sequence.

**Restriction enzyme cloning**

Choosing restriction sites:

1. Check the sequence of your POI DNA template for BamHI, NheI, AgeI, and XbaI recognition sites and choose 2 sites in the MCS that do not occur in your template.
2. If possible, choose the outermost sites to avoid adding excess amino acids to the N- or C-terminus of your POI (see **Fig. 1**).
3. Avoid using both NheI and XbaI when possible because these two enzymes create matching overhangs. This thus reduces the chance of successful ligation in the correct orientation.

Designing primers:

1. Make sure that you are using the correct reading frame of the template DNA and that it does not contain premature stop codons.
2. Design both forward and reverse primers to be between 20-40 base pairs (bp) in length and to have a melting temperature ($T_M$) between 58-64 C°. Make sure each primer ends in a G or C at the 3' end.
3. The 5' end of each primer must contain a string of 6 nucleotides (of any sequence) followed by the restriction site. This is needed to provide a toehold for the restriction enzyme.
4. Design the reverse primer to be reverse complement of the template DNA sequence.
5. Check primers for both intra- and intermolecular high $T_M$ hairpins using e.g. IDT's oligoanalyzer (https://www.idtdna.com/pages/tools/oligoanalyzer) and introduce changes to remove these if necessary.

Run PCR reaction:

1. Mix reaction components in a 0.2 ml PCR tube on ice

**PCR reaction (50 µl)**

| **Volume (µl)** | **Reagent** |
|---|---|

| | |
|---|---|
| 25 | Q5 High-Fidelity 2x Master Mix (NEB, USA) |
| 1 | 10 ng/µl template |
| 0.5 | 50 µM forward primer |
| 0.5 | 50 µM reverse primer |
| 23 | dd$H_2O$ |

2. Program thermocycler with the following settings and run PCR:

**Thermocycler settings**

| # | Step | Temperature | Time | |
|---|---|---|---|---|
| 1 | Initial denaturation | 98°C | 30 sec | |
| 2 | Denaturation | 98°C | 10 sec | 30x cycles of steps 2-4 |
| 3 | Annealing | 58-64°C | 30 sec | |
| 4 | Extension | 72°C | 30 sec per 1 kbp | |
| 5 | Hold | 4-10°C | ∞ | |

kbp = kilo base pair

3. Purify reactions using QIAquick PCR Purification kit (QIAGEN, Netherlands)
4. Elute in 42 µl dd$H_2O$

Double restriction enzyme digest of purified PCR product:

1. Set up restriction digest reactions of purified PCR product and plasmid:

**PCR product digest reaction (50 µl)**

| Volume (µl) | Reagent |
|---|---|
| 5 | 10x CutSmart buffer (NEB, USA) |
| 1.5 | Restriction enzyme 1* (30 Units) |
| 1.5 | Restriction enzyme 2* (30 Units) |
| 42 | Purified PCR product |

* Use of NEB high fidelity (HF) enzymes is recommended

**Plasmid digest reaction (50 µl)**

| Volume (µl) | Reagent |
|---|---|
| 5 | 10x CutSmart buffer (NEB, USA) |
| 1.5 | Restriction enzyme 1* (30 Units) |
| 1.5 | Restriction enzyme 2* (30 Units) |
| X | 5 µg plasmid DNA |
| Add to 50 | dd$H_2O$ |

2. Mix well and incubate digests at 37˚C for 1.5 h
3. Add 2 µl of 1U/µl FastAP Alkaline phosphatase (Thermo Fisher Scientific, USA) **only** to the plasmid digest to remove 5’ phosphates (prevents self-ligation of insert-less plasmid)
4. Incubate both digests for another 30 min at 37˚C
5. Mix digests with 10 µl Gel Loading Dye, Purple (6x) (NEB, USA)
6. Run on a 1% (w/v) agarose gel made up in 1x TAE buffer and supplemented with 1x SYBR Safe DNA Gel Stain (Thermo Fisher Scientific, USA) for 30 min at 150 V
7. Excise bands with clean razor blades
8. Purify excised DNA bands with Zymoclean Gel DNA Recovery Kit (Zymo Research, USA)
9. Elute in two steps with 2x 10 µl dd$H_2O$ (20 µl final volume)
10. Measure DNA concentration using a NanoDrop spectrophotometer (Thermo Fisher Scientific, USA)

Ligation

1. Using the size in bp of both digested insert and plasmid fragments, calculate the amount of insert in ng to ligate with 100 ng dephosphorylated plasmid using a 2:1 insert:plasmid ratio → Use the NEBioCalculator: https://nebiocalculator.neb.com/#!/ligation
2. Set up the ligation reaction and include a negative control reaction in which the insert is replaced with $ddH_2O$

**Ligation reaction (10 µl)**

| Volume (µl) | Reagent |
|---|---|
| 5 | 2x Quick Ligase Reaction Buffer |
| X | X ng of insert DNA |
| X | 100 ng of plasmid DNA |
| 0.5 | Quick ligase (NEB, USA) |
| Add to 10 | $ddH_2O$ |

3. Mix well and incubate for 15 min at room temperature

Transformation

1. Thaw one 100 µl vial of chemically competent *E. coli* Stellar cells (Takara Bio, Japan) per two ligation reactions for 10 min on ice
2. Split into 2x 50 µl aliquots in two separate tubes on ice and add 5 µl of ligation reaction to each
3. Incubate for 30 min on ice
4. Heat shock tubes at 42˚C for 35 sec in a heat block or water bath
5. Quickly remove to ice and chill for 1-2 min
6. Rescue by addition of 200 µl SOC recovery medium
7. Incubate tubes at 37˚C for 30-60 min shaking at 1,200 rpm
8. Plate out ˜150 µl on LB-Carb agar plates, let dry and incubate upside down overnight at 37˚C

**Gibson cloning**

Primer design

Use NEBuilder Assembly tool to design insert primers with matching overhangs for Gibson assembly: https://nebuilder.neb.com/#!/

1. Generate a new fragment and copy&paste backbone plasmid DNA sequence, click process text and check the 'vector' and 'circular' boxes. Rename 'new fragment' to 'plasmid'.
2. Select 'restriction digest' as the method for production of a linearized fragment and specify your choice of restriction sites. Finally click 'Add'.
3. Generate another new fragment and copy&paste insert template DNA sequence, click process text and check the 'vector' and 'circular' boxes if applicable. Rename to 'insert'.
4. Select 'PCR' as the method for production of a linearized fragment and specify the start and end base of your insert. Include stop codon if needed. Finally click 'Add'.
5. The exonuclease in the Gibson assembly mix will remove the 4 bp overhangs generated by restriction digest of the plasmid, leaving only 1 base of the original 6 bp recognition site behind. In order to maintain a proper open reading frame, upstream and/or downstream junctions between plasmid and insert fragments may need to be adjusted by adding either two or five bases, to restore a single codon or the complete restriction site, respectively. Click on the pencil symbol of the newly added insert fragment to edit these junctions.
6. The program generates an 'assembled sequence' that should be thoroughly inspected to contain the insert at the correct location and in the correct reading frame.

7. If all looks well order the suggested insert primer pair containing the proper Gibson assembly overhangs.

PCR amplification of insert fragment

1. Set up and run PCRs as described above

Restriction digest to create plasmid fragment

1. While the insert PCR is running, set up the plasmid restriction digest reaction as described above and include the phosphatase treatment step

Gel purification

1. Mix insert PCR and plasmid restriction digest reaction with 10 µl Gel Loading Dye, Purple (6x) (NEB, USA), run on a 1% (w/v) agarose and purify from excised gel band as described above

Gibson assembly reaction

1. Calculate the volume of insert and vector fragment that contains 50 fmoles of each using NEBioCalculator: https://nebiocalculator.neb.com/#!/dsdnaamt
2. Mix calculated volume of both vector and insert fragment and then dilute two-fold with 2x Gibson Assembly Master Mix (NEB,USA)
3. Incubate at 50˚C for 30 min
4. Transform up to 5 µl into chemical competent *E. coli* Stellar cells via heat shock as described above
